# Common Fronto-temporal Effective Connectivity in Humans and Monkeys

**DOI:** 10.1101/2020.04.03.024042

**Authors:** Francesca Rocchi, Hiroyuki Oya, Fabien Balezeau, Alexander J. Billig, Zsuzsanna Kocsis, Rick Jenison, Kirill V. Nourski, Christopher K. Kovach, Mitchell Steinschneider, Yukiko Kikuchi, Ariane E. Rhone, Brian J. Dlouhy, Hiroto Kawasaki, Ralph Adolphs, Jeremy D.W. Greenlee, Timothy D. Griffiths, Matthew A. Howard, Christopher I. Petkov

## Abstract

Cognitive pathways supporting human language and declarative memory are thought to have uniquely evolutionarily differentiated in our species. However, cross-species comparisons are missing on site-specific effective connectivity between regions important for cognition. We harnessed a new approach using functional imaging to visualize the impact of direct electrical brain stimulation in human neurosurgery patients. Applying the same approach with macaque monkeys, we found remarkably comparable patterns of effective connectivity between auditory cortex and ventro-lateral prefrontal cortex (vlPFC) and parahippocampal cortex in both species. Moreover, in humans electrical tractography revealed rapid evoked potentials in vlPFC from stimulating auditory cortex and speech sounds drove vlPFC, consistent with prior evidence in monkeys of direct projections from auditory cortex to vocalization responsive regions in vlPFC. The results identify a common effective connectivity signature that from auditory cortex is equally direct to vlPFC and indirect to the hippocampus (via parahippocampal cortex) in human and nonhuman primates.

**Highlights:**

- Privileged human auditory to inferior frontal connectivity, linked to monkeys
- Common auditory to parahippocampal effective connectivity in both species
- Greater lateralization in human effective connectivity, more symmetrical in monkeys
- Human fronto-temporal network function rooted in evolutionarily conserved signature

**eTOC short summary:** Functional connectivity between regions crucial for language and declarative memory is thought to have substantially differentiated in humans. Using a new technique to similarly visualize directional effective connectivity in humans and monkeys, we found remarkably comparable connectivity patterns in both species between fronto-temporal regions crucial for cognition.

## INTRODUCTION

Brain networks adapted for specialized functions typically show direct, rapid or effective connectivity between regions crucial for behavior. Finding such connectivity, alongside evidence for evolutionary homology, convergence or divergence, can be of substantial theoretical significance: Within the motor domain, human and nonhuman primates have direct cortico-spinal projections subserving fine movement control that are indirect in rodents^1^. Also, human laryngeal motor cortex projects directly to a brain stem nucleus (ambiguus) controlling laryngeal muscles^2^. Such projections for vocal production are more indirect in nonhuman primates^3^ and rodents^4^, shedding light on human speech evolution^5^.

Language defines us as a species and because of its prominent role in declarative memory substantial evolutionary differentiation of human cognitive pathways is expected. Comparative studies often see considerable levels of evolutionary conservation alongside insights on species-specific differences^6-12^. Yet, certain cross-species comparisons are missing, such as on the impact of directed effective connectivity with the required precision of site-specific perturbation that can be applied to both human and nonhuman primates. Thereby, the question on the extent of differentiation versus conservation in primate fronto-temporal systems—although crucial for understanding which aspects of human language and cognition can find realistic nonhuman animal models—remains open.

Speech and language are supported by a fronto-temporal network, including auditory and ventro-lateral prefrontal cortex (vlPFC) areas interconnected via white matter pathways^9^. Left hemisphere areas posterior to Heschl’s gyrus (HG; an anatomical landmark associated with primary auditory cortex) are interconnected with Brodmann areas 44 and 45 in vlPFC by way of the dorsal arcuate fasciculus pathway^13^. There is now evidence for an auditory homolog in chimpanzees and macaques, with its left hemisphere lateralization as a prominent human-specific difference^8^. In monkeys, vlPFC neurons respond to vocalization sounds^14^, and single neuron tractography has shown evidence for directional connectivity between non-primary auditory (lateral belt) areas and vlPFC^15^. Whether similar auditory to inferior frontal interconnectivity exists in humans is however unclear^7,16,17^. Based on the notion of human vlPFC areas 44 and 45 having functionally differentiated for language^9^, one prediction is that human auditory cortex would have a greater effective connectivity impact on areas 44/45. By contrast, the impact on the adjacent frontal operculum (FOP) could be similar in humans and monkeys^18^.

Sensory input to the medial temporal lobe (MTL) in humans is important for declarative memory and may also have differentiated in humans. Monkey recognition memory for sounds is surprisingly more fleeting than for visual items^19^. Also, studies in nonhuman animals (rodents, cats and monkeys) show primarily indirect projections from auditory cortex to the hippocampus^20^ via parahippocampal/perirhinal cortex^21,22^. However, human neuroimaging studies have not been able to demonstrate direct auditory to hippocampal connectivity, with the current data suggesting indirect interconnectivity with auditory cortex (via parahippocampal cortex)^23^, as seen in other species. Thus, another prediction, given that human language allows naming and conceptualizing sounds and thereby better remembering them^24^, is that the auditory-to-MTL memory circuit may be largely evolutionarily conserved.

New approaches for assessing effective connectivity similarly across the species could shed light on human cognitive pathway specialization or primate origins. In monkeys, electrical stimulation combined with functional magnetic resonance imaging (esfMRI) was developed to visualize the impact of site-specific stimulation on regional effective connectivity^25^. Direct electrical brain stimulation is a common treatment for debilitating brain disorders, thus following safety testing the esfMRI method was recently translated to human patients being monitored for neurosurgery^26^. We conducted a comparative study of esfMRI effects obtained by stimulating auditory cortex in humans and macaques. Electrical stimulation of auditory cortex induced differential esfMRI activity in several vlPFC (areas 44, 45 and FOP) and MTL subregions (including parahippocampal and hippocampal areas). We found largely comparable effective connectivity patterns across the species. In humans, we also studied electrical stimulation tractography to assess the latency of interconnectivity between auditory cortex, vlPFC and MTL. Finally, we observed strong neurophysiological responses to speech sounds in human vlPFC and establish directional effective interconnectivity with auditory cortex.

## RESULTS

### Auditory cortex esfMRI in macaques

We first studied the esfMRI response from stimulation of auditory cortical sites in the right hemisphere of two macaques. Auditory cortex esfMRI induced significantly stronger activity relative to non-stimulation trials (Fig. 1; *p* < 0.05, cluster corrected, *Z* > 2.8) in areas including auditory cortex, prefrontal and MTL regions (Table 1). The prefrontal pattern involving area 8 and areas 44/45 is generally similar to prefrontal neuronal tractography from auditory lateral belt in macaques^15,27^ (Table 1). MTL activation included parahippocampal gyrus^20^.

**Figure 1.**
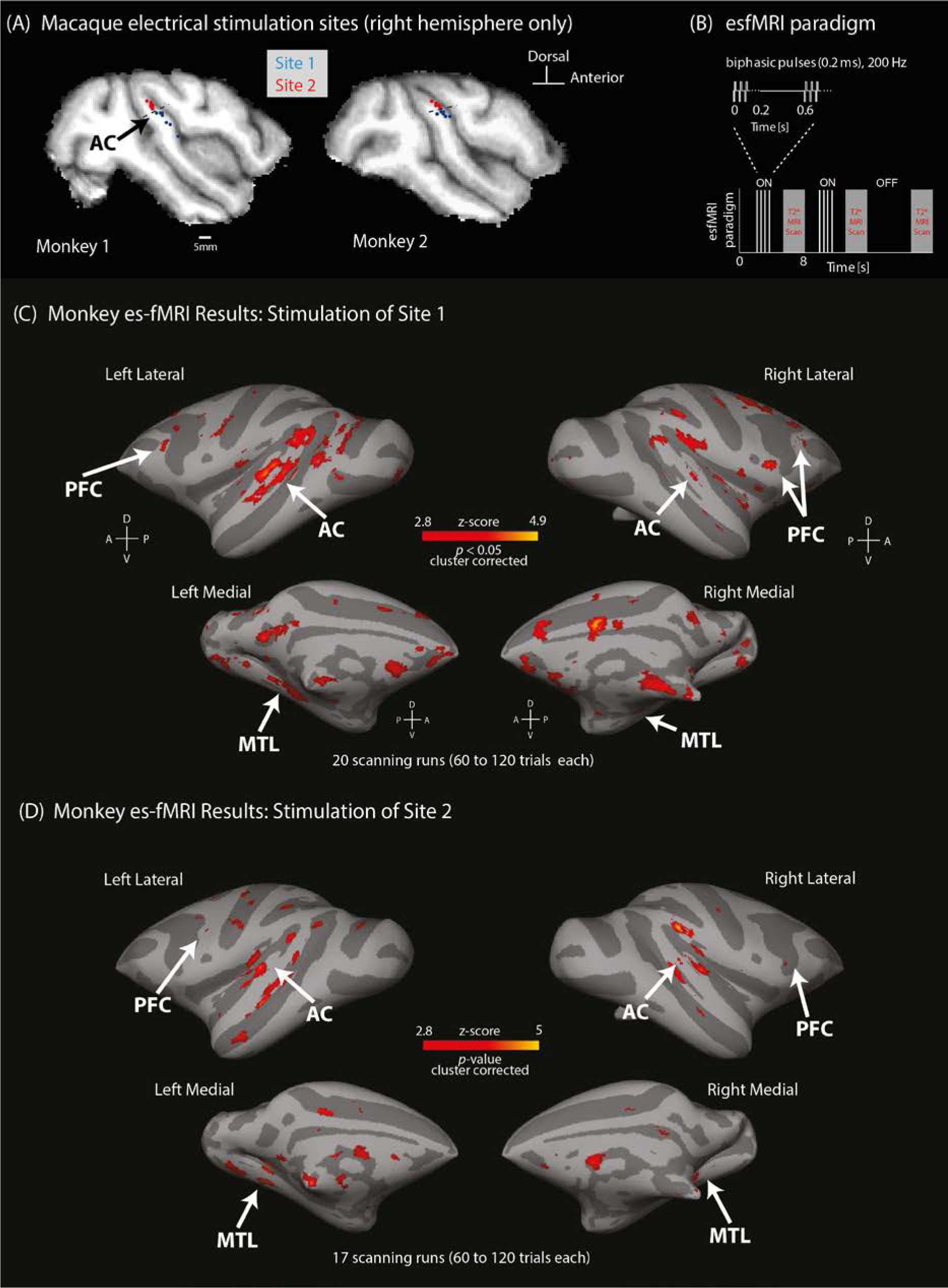
Macaque monkey auditory cortex electrical stimulation sites and esfMRI results. (A) Stimulation sites 1 (blue) and 2 (red) auditory cortex of the right hemisphere in the two macaques (M1 and M2). (B) esfMRI paradigm timing. (C-D) Macaque esfMRI group results showing significantly activated voxels relative to no-stimulation trials in response to electrical stimulation of the auditory cortex sites: Site 1 (C) and Site 2 (D), cluster-corrected *p* < 0.05, *Z* > 2.8 (see Table 1 for list of activated anatomical regions). Results projected to the surface-rendered standard macaque template brain. Abbreviations: auditory cortex (AC), prefrontal cortex (PFC) and medial temporal lobe (MTL).

**Table 1.** Macaque esfMRI results as significant activation in the listed anatomical regions. Shown are the macaque brain regions in reference to the macaque atlas^61^ (x,y,z stereotactic coordinates), the peak voxel Z-value in the region and the number of voxels above *Z* > 2.8 corrected significance threshold. Right (rt.) and left (lt.) hemispheres are identified.

Site 1 esfMRI macaque group results
| size (voxels) | peak z-score | x y z | *Temporal-Occipital Cortex* |
| --- | --- | --- | --- |
| 1350 | 3.89 | 6 -13 17 | rt. Primary/Secondary Visual Cortex (e.g., V2, V1) |
| 1296 | 4.19 | -4 -6 21 | lt. Primary/Secondary Visual Cortex (e.g., V2, V1) |
| 268 | 3.93 | -24 6 20 | lt. Superior Temporal Sulcus STS, dorsal (TAa) |
| 185 | 3.70 | -24 3 20 | lt. Inferior Temporal Cortex (TEO) |
| 118 | 4.10 | -24 9 18 | lt. Koniocortex (Primary Auditory) |
| 86 | 3.29 | 22 9 23 | rt. Koniocortex (Primary Auditory) |
| 86 | 3.30 | -14 -11 29 | lt. Higher Visual Cortex (V3a) |
| 77 | 3.30 | 23 13 17 | rt. Auditory Core, field R |
| 61 | 3.38 | 15 2 7 | rt. Higher Visual Cortex (V4v) |
| 55 | 3.74 | 10 0 15 | rt. Medial Occipital area (V2) |
| 46 | 3.19 | -21 -3 19 | lt. Fundus of Superior Temporal Sulcus (FST) |
| 44 | 3.48 | 27 11 19 | rt. Auditory Non-primary Cortex |
| 39 | 3.43 | -12 3 7 | lt. Higher Visual Cortex (V4v) |
| 36 | 3.60 | -22 10 16 | lt. Auditory Core, field R |
| 33 | 3.09 | -15 -11 27 | lt. Higher Visual Area, dorsal (V3d) |
| 31 | 3.36 | -27 12 13 | lt. Rostral Auditory Para-Belt (RPB) |
| 21 | 3.08 | -18 7 13 | lt. STS fundus (IPa) |
| 21 | 2.98 | 3 -12 31 | rt. Higher Visual Cortex (V6av) |
| 18 | 3.37 | -5 -11 27 | lt. Higher Visual Area (V6) |
| 16 | 3.01 | 1 -12 25 | rt. Higher Visual Area (V6) |
| 11 | 3.15 | 17 17 6 | rt. STS fundus (IPa) |
| size (voxels) | peak z-score | x y z | *Frontal cortex* |
| 699 | 4.02 | 12 23 33 | rt. Frontal Eye Fields (area 8av) |
| 369 | 4.08 | 1 28 32 | rt. Supplementary Motor Area (SMA) |
| 349 | 3.55 | -9 -4 12 | lt. Area 10 |
| 298 | 4.00 | 5 32 23 | rt. Area 11 |
| 291 | 3.92 | 21 22 10 | rt. Frontal-Operculum |
| 270 | 3.77 | -6 31 32 | lt. Frontal Eye Fields (area 8av) |
| 252 | 3.63 | 11 1 10 | rt. Area 10 |
| 238 | 3.37 | -21 20 25 | lt. Primary Motor Cortex (4/F1) |
| 214 | 3.97 | 2 33 12 | rt. Medial Prefrontal Cortex (10m) |
| 167 | 3.74 | 8 33 26 | rt. Dorso-Lateral Prefrontal Cortex (8b) |
| 150 | 3.65 | -3 34 13 | lt. Medial Prefrontal Cortex (10m) |
| 145 | 3.70 | 13 16 30 | rt. Primary Motor Cortex (4/F1) |
| 129 | 3.59 | -19 29 23 | lt. Frontal Eye Fields (area 8av) |
| 119 | 3.57 | 2 38 30 | rt. Dorso-Lateral Prefrontal Cortex (8bd) |
| 114 | 3.39 | -6 32 31 | lt. Premotor Area, dorsal 6dr |
| 112 | 3.80 | 4 34 12 | rt. Orbito-frontal (13l) |
| 108 | 3.37 | -20 16 16 | lt. Frontal-Operculum |
| 85 | 3.59 | 2 37 31 | rt. Premotor Area, dorsal 6dr |
| 63 | 3.49 | -1 29 32 | lt. Supplementary Motor Area (SMA) |
| 57 | 3.21 | -4 37 15 | lt. Orbito-prefrontal (13m) |
| 48 | 3.49 | 24 27 16 | rt. Premotor Area, ventral (6Va) |
| 44 | 3.26 | 20 20 8 | rt. Area 46 |
| 31 | 3.44 | 13 33 15 | rt. Orbito-prefrontal (13m) |
| 27 | 3.19 | -15 26 25 | lt. Premotor Area, ventral (6Va) |
| 27 | 3.32 | -19 29 22 | lt. Ventro-Lateral Prefrontal Cortex (45b) |
| 26 | 3.23 | -14 26 24 | lt. Dorso-Lateral Prefrontal Cortex (8b) |
| 16 | 3.12 | -10 33 12 | lt. Orbito-frontal (13l) |
| 15 | 3.05 | 15 25 7 | rt. Agranular Insular, lateral (Ial) |
| 15 | 3.34 | 13 31 18 | rt. Area 44 |
| 10 | 2.98 | 14 29 19 | rt. Area 45 |
| 7 | 3.13 | -19 29 22 | lt. Area 45 |

| **size (voxels)** | **peak z-score** | **x y z** | ***Medial Temporal Lobe*** |
| --- | --- | --- | --- |
| 129 | 3.49 | 15 -1 13 | rt. Hippocampus (CA1) |
| 118 | 3.58 | -9 6 7 | lt. Parahippocampal Cortex (TFO) |
| 90 | 3.50 | -10 6 8 | lt. Parahippocampal Cortex (TH) |
| 73 | 3.39 | 9 6 8 | rt. Parahippocampal Cortex (TFO) |
| 59 | 3.45 | 16 1 13 | rt. Hippocampus (CA3) |
| 52 | 3.44 | 12 2 12 | rt. Subiculum |
| 44 | 3.52 | 10 7 6 | rt. Parahippocampal Cortex (TH) |
| 40 | 3.16 | 16 8 7 | rt. Dentate Gyrus |
| 38 | 3.60 | 11 3 12 | rt. Prosubiculum |
| 36 | 3.54 | -16 8 6 | lt. Dentate Gyrus |
| 35 | 3.68 | -17 9 6 | lt. Hippocampus (CA3) |
| 34 | 3.16 | 15 8 7 | rt. Hippocampus (CA4) |
| 25 | 3.37 | -13 3 8 | lt. Hippocampus (CA1) |
| 25 | 3.23 | -12 5 8 | lt. Subiculum |
| 22 | 3.56 | 16 0 14 | rt. Hippocampus (CA2) |
| 21 | 3.38 | -12 3 9 | lt. Prosubiculum |
| 14 | 3.23 | -15 9 6 | lt. Hippocampus (CA4) |
| 10 | 3.42 | -17 9 6 | lt. Hippocampus (CA2) |
| 10 | 3.14 | -9 7 6 | lt. Parahippocampal Cortex (TF) |

| **size (voxels)** | **peak z-score** | **x y z** | ***Cingulate Cortex*** |
| --- | --- | --- | --- |
| 460 | 4.94 | 4 19 28 | rt. Anterior Cingulate Cortex (24c) |
| 145 | 3.62 | 3 14 27 | rt. Posterior Cingulate Cortex (23c) |
| 94 | 3.48 | 1 2 26 | rt. Posterior Cingulate Cortex (23b) |
| 90 | 3.82 | 1 34 21 | rt. Anterior Cingulate Cortex (24a) |
| 83 | 4.25 | 2 18 27 | rt. Anterior Cingulate Cortex (24b) |
| 38 | 3.64 | -1 35 20 | lt. Anterior Cingulate Cortex (24a) |
| 29 | 3.12 | -6 24 30 | lt. Anterior Cingulate Cortex (24c) |
| 11 | 2.97 | -1 3 26 | lt. Posterior Cingulate Cortex (23b) |
| 11 | 3.02 | -3 -1 26 | lt. Posterior Cingulate Cortex (31) |
| 10 | 3.20 | -2 -6 22 | lt. Posterior Cingulate Cortex (v23b) |

| **size (voxels)** | **peak z-score** | **x y z** | ***Parietal Cortex*** |
| --- | --- | --- | --- |
| 384 | 3.70 | -21 15 19 | lt. Somatosensory Area secondary (SII) |
| 315 | 3.55 | 22 10 23 | rt. Somatosensory Area secondary (SII) |
| 189 | 3.29 | 4 -8 27 | rt. Medial Parietal Area (7m/PGm) |
| 143 | 3.75 | -6 -10 28 | lt. Medial Intraparietal Area (MIP) |
| 124 | 3.17 | 16 4 35 | rt. Intraparietal Area (LIPd) |
| 56 | 3.09 | 26 14 17 | rt. Somatosensory Cortex (areas 1-2) |
| 14 | 3.00 | 7 -11 28 | rt. Medial Intraparietal Area (MIP) |
| 10 | 3.05 | -11 -7 33 | lt. Inferior Parietal Lobule, posterior (7a/Opt/PG) |

| **size (voxels)** | **peak z-score** | **x y z** | ***Other Areas*** |
| --- | --- | --- | --- |
| 1148 | 4.11 | 9 16 22 | rt. Striatum |
| 843 | 4.05 | -2 25 14 | lt. Striatum |
| 570 | 4.03 | 21 21 10 | rt. Thalamus |
| 286 | 3.95 | 18 13 14 | rt. Claustrum |
| 229 | 3.69 | -21 15 17 | lt. Granular Insular Cortex |
| 227 | 3.67 | 5 7 15 | rt. Superior Colliculus |
| 208 | 3.40 | 19 13 16 | rt. Granular Insular Cortex |
| 181 | 3.73 | 4 5 11 | rt. Inferior Colliculus |
| 137 | 3.93 | 8 4 18 | rt. Medial Pulvinar |
| 67 | 3.40 | -4 5 9 | lt. Inferior Colliculus |
| 42 | 3.44 | 10 4 19 | rt. Lateral Pulvinar |
| 29 | 3.73 | 15 14 12 | rt. Globus Pallidus (GPe/Gpi) |
| 28 | 3.03 | -4 5 10 | lt. Superior Colliculus |
| 27 | 3.28 | 11 5 12 | rt. Inferior Pulvinar |
| 21 | 3.05 | -17 13 12 | lt. Claustrum |
| 13 | 3.04 | -8 3 14 | lt. Lateral Pulvinar |
| 11 | 3.21 | 8 16 3 | rt. Nucleus Accumbens |
| 10 | 3.15 | 4 29 9 | rt. Anterior Olfactory Nucleus<br>(AONd/m) |
| 10 | 2.93 | 7 20 6 | rt. Nucleus of Lateral Olfactory Tract<br>(NLOT) |

**Site 2 esfMRI macaque group results**
| **size (voxels)** | **peak z-score** | **x y z** | ***Temporal-Occipital Cortex*** |
| --- | --- | --- | --- |
| 652 | 4.16 | -14 -5 18 | lt. Secondary Visual Cortex (V2) |
| 155 | 3.96 | 21 10 19 | rt. Koniocortex (Primary Auditory) |
| 154 | 3.89 | 12 -1 13 | rt. Secondary Visual Cortex (V2) |
| 132 | 4.32 | 27 6 22 | rt. Superior Temporal Sulcus STS, dorsal<br>(TAa) |
| 66 | 3.86 | 27 8 23 | rt. Auditory Non-primary Cortex |
| 57 | 3.30 | -7 0 17 | lt. Medial Occipital area (V2) |
| 56 | 3.58 | -18 15 3 | lt. STS fundus (IPa) |
| 30 | 3.10 | 18 11 7 | rt. STS fundus (IPa) |
| 29 | 3.65 | 22 11 17 | rt. Auditory Core, field R |
| 20 | 3.36 | -24 9 14 | lt. Superior Temporal Sulcus STS, dorsal (TAa) |
| 14 | 3.51 | -22 9 17 | lt. Koniocortex (Primary Auditory) |
| 13 | 3.53 | -14 -5 11 | lt. Higher Visual Cortex (V4v) |
| 13 | 3.36 | 23 -3 29 | rt. Higher Visual Cortex (V4t) |

| **size (voxels)** | **peak z-score** | **x y z** | ***Frontal cortex*** |
| --- | --- | --- | --- |
| 523 | 4.30 | 23 2 28 | rt. Area 11 |
| 263 | 3.98 | -14 -7 11 | lt. Area 10 |
| 114 | 3.68 | -17 19 25 | lt. Primary Motor Cortex (4/F1) |
| 29 | 3.18 | -11 12 32 | rt. Area 46 |
| 15 | 3.20 | -18 15 16 | lt. Frontal_Operculum |
| 7 | 3.10 | 21 15 16 | rt. Frontal_Operculum |
| 3 | 2.99 | 19 29 16 | rt. Area 44 |

| size (voxels) | peak z-score | x y z | *Medial Temporal Lobe* |
| --- | --- | --- | --- |
| 88 | 3.51 | -7 3 11 | lt. Parahippocampal Cortex (TFO) |
| 38 | 3.75 | -14 -1 15 | rt. Hippocampus (CA3) |
| 26 | 3.48 | 8 3 12 | rt. Parahippocampal Cortex (TFO) |
| 14 | 3.34 | 13 -1 14 | rt. Subiculum |
| 13 | 3.54 | -10 3 12 | lt. Prosubiculum |
| 11 | 3.24 | -14 -2 13 | lt. Hippocampus (CA1) |

| size (voxels) | peak z-score | x y z | *Parietal Cortex* |
| --- | --- | --- | --- |
| 74 | 3.55 | -18 13 22 | lt. Somatosensory Area secondary (SII) |
| 73 | 3.19 | 23 12 18 | rt. Somatosensory Area secondary (SII) |
| 42 | 3.40 | 22 -3 30 | rt. Inferior Parietal Lobule, posterior<br>(7a/Opt/PG) |

| size (voxels) | peak z-score | x y z | *Other Areas* |
| --- | --- | --- | --- |
| 408 | 3.96 | 4 29 18 | rt. Striatum |
| 304 | 3.97 | 20 12 17 | rt. Granular Insular Cortex |
| 295 | 4.38 | -2 5 11 | lt. Inferior Colliculus |
| 226 | 3.87 | -2 26 18 | lt. Striatum |
| 199 | 3.73 | -4 5 10 | lt. Superior Colliculus |
| 197 | 4.64 | 27 6 23 | rt. Thalamus |
| 157 | 3.51 | -17 11 19 | lt. Granular Insular Cortex |
| 58 | 3.31 | -9 3 14 | lt. Lateral Pulvinar |
| 45 | 3.70 | -6 7 12 | lt. Medial Geniculate, magnocellular |
| 44 | 3.20 | -18 15 16 | lt. Claustrum |
| 28 | 3.36 | 14 25 11 | rt. Claustrum |
| 26 | 3.31 | -7 7 12 | lt. Medial Geniculate, ventral |
| 16 | 3.47 | -10 4 13 | lt. Inferior Pulvinar |
| 12 | 3.09 | 7 3 12 | rt. Inferior Colliculus |
| 10 | 3.04 | -13 15 16 | lt. Globus Pallidus (GPe/Gpi) |

Grouping of stimulation sites was conducted across two tonotopically organized fields (‘core’ Site 1 and ‘belt’ Site 2; Suppl. Fig. 1). However, esfMRI results from stimulating these sites were statistically indistinguishable (no significant voxels surviving the *p* < 0.05 cluster-corrected threshold for the Site 1 versus Site 2 contrast, or vice versa). We suspected that this might arise because of partially overlapping stimulation, which we confirmed by estimating the passive current spread using the formula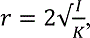
, where *I* is current in mA, and *K* a constant of pyramidal cell excitability for a 200 µs pulse (1.3 mA/mm^2^). The resulting radius (*r*) of passive current spread is 0.9 mm, confirming that stimulating core auditory cortex sites also likely passively electrically stimulates portions of the adjoining auditory belt, and vice versa, an important consideration for interpreting the esfMRI results.

### Auditory cortex esfMRI in humans

We assessed the esfMRI activity in response to human auditory cortex electrical stimulation of depth electrodes at medial or lateral sites in either left or right Heschl’s gyrus (HG; Fig. 2A; Suppl. Fig. S2 shows individual subject contact locations). Stimulation sites were subdivided into Site 1 (medHG), grouping contacts in medial HG areas where significant phase-locking to high-repetition click sound rates occur (Methods), and Site 2, grouping lateral HG and planum temporale contacts (latHG + PT) lacking high click rate neural phase locking (Suppl. Fig. S3). Suppl. Fig. S4 shows the amount of data retained following removal of neurosurgically resected sites and loss of data around electrode contacts.

**Figure 2.**
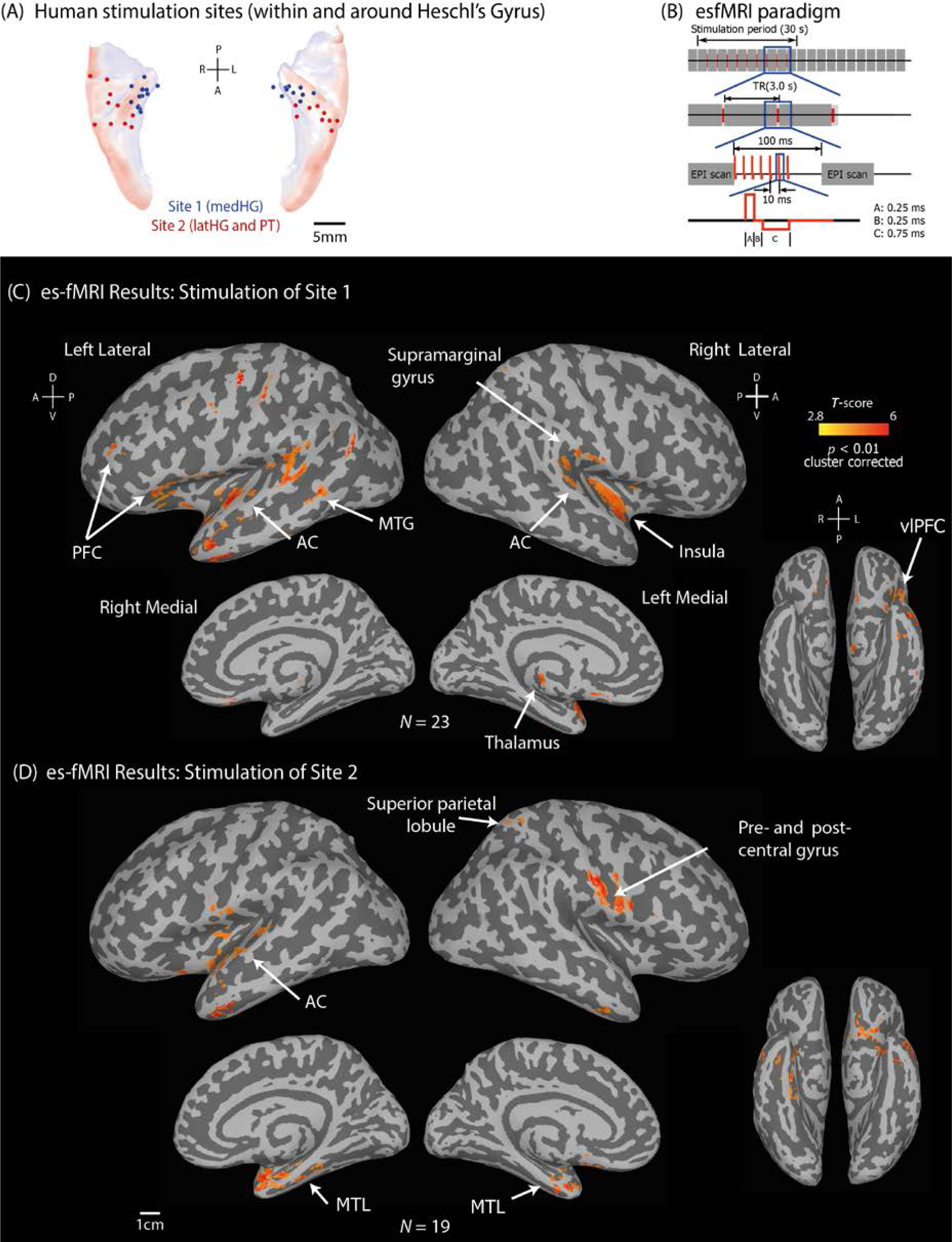
Human auditory cortex electrical stimulation sites and esfMRI results. (A) Stimulation of depth electrodes in the transverse temporal gyrus (Heschl’s gyrus; HG). Human auditory stimulation Sites 1 and 2 shown looking down on the superior temporal plane. Stimulation sites are identified at the center of the adjacent contacts used for stimulation (Suppl. Fig. S2 shows actual contact location in each participant). (B) esfMRI paradigm timing. (C-D) Human esfMRI group results shown as significantly activated voxels relative to no-stimulation trials (cluster corrected, *T* = 2.8, *p* < 0.01) in response to electrical stimulation of Site 1 (C) or Site 2 (D), shown on the surface-rendered Montreal Neurological Institute human standard brain.

Significant esfMRI effects (Fig. 2C; cluster corrected *p* < 0.05) from stimulating Site 1 (medHG) included auditory cortex (superior temporal gyrus, STG) and vlPFC (inferior frontal gyrus; Table 2). Site 2 (latHG+PT) stimulation activated areas including STG and MTL (Fig. 2D). In some individual participants, the activity response was significant in vlPFC (Suppl. Fig. S5) and MTL (Suppl. Fig. S6). The strength of the esfMRI response in vlPFC and MTL overlaid on the stimulated HG contacts that produced it is shown in Suppl. Fig. S7.

**Table 2.** Human esfMRI results, as significant activation in the listed anatomical regions. Shown are the human brain regions in reference to the human atlas^67^ in MNI x,y,z coordinates, the peak voxel value in the region and the number of voxels above the corrected significance threshold (cluster-wise alpha < 0.5 with primary *p* < 0.005).

**Site 1 (medHG) esfMRI human group results:**
| size (voxel) | peak T | MNI<br>coordinates<br>(mm) |  |  | Brain structure |
| --- | --- | --- | --- | --- | --- |
|  |  | x | y | z |  |
| 831 | 7.38 | 54.5 | 0.5 | -28.5 | lt. middle temporal gyrus |
| 327 | 7.10 | 66.5 | 56.5 | 13.5 | lt. superior temporal gyrus |
| 291 | 5.77 | -41.5 | 4.5 | -12.5 | rt. insula |
| 241 | 5.62 | -65.5 | 26.5 | 27.5 | rt. supramarginal gyrus |
| 166 | 4.97 | 42.5 | -29.5 | 1.5 | lt. inferior frontal gyrus |
| 149 | 6.54 | 2.5 | 26.5 | -4.5 | lt. thalamus |
| 123 | 4.57 | -32.5 | -18.5 | -2.5 | rt. putamen |
| 84 | 5.26 | 32.5 | -47.5 | 21.5 | lt. middle frontal gyrus |

**Site 2 (latHG + PT) esfMRI human group results:**
| size (voxel) | peak T | MNI<br>coordinates<br>(mm) |  |  | Brain structure |
| --- | --- | --- | --- | --- | --- |
|  |  | x | y | z |  |
| 696 | 7.29 | -23.5 | -5.5 | -26.5 | rt. parahippocampal gyrus |
| 600 | 7.77 | 46.5 | 2.5 | -30.5 | lt. inferior temporal gyrus |
| 377 | 7.19 | -63.5 | 10.5 | 37.5 | rt. postcentral gyrus |
| 173 | 7.34 | 58.5 | -11.5 | -6.5 | lt. superior temporal gyrus |
| 140 | 7.58 | -31.5 | 64.5 | 65.5 | rt. superior parietal lobule |
| 104 | 4.68 | -15.5 | 21.5 | -17.5 | lt. superior orbital gyrus |

**Site 1 versus Site 2 esfMRI human group results:**
| size (voxel) | peak T | MNI<br>coordinates<br>(mm) |  |  | Brain structure |
| --- | --- | --- | --- | --- | --- |
|  |  | x | y | z |  |
| 430 | 7.21 | 66.5 | 56.5 | 13.5 | lt. middle and superior temporal gyrus |
| 196 | 7.17 | -63.5 | 28.5 | 25.5 | rt. supramarginal gyrus |
| 106 | 5.39 | 48.5 | -15.5 | 1.5 | lt. inferior frontal gyrus |

**Site 2 versus Site 1 esfMRI human group results:**
| size (voxel) | peak T | MNI<br>coordinates<br>(mm) |  |  | Brain structure |
| --- | --- | --- | --- | --- | --- |
|  |  | x | y | z |  |
| 971 | 5.65 | -35.5 | -19.5 | -28.5 | rt. ATL (incl. entorhinal cortex,<br>parahippocampal gyrus, temporal<br>pole & amygdala) |
| 283 | 5.60 | 22.5 | -5.5 | -28.5 | lt. ATL (incl. entorhinal cortex &<br>amygdala) |
| 262 | 5.71 | -63.5 | 6.5 | 21.5 | lt. precentral gyrus |

Unlike the monkey esfMRI results, which did not significantly differ between the two auditory cortex stimulation sites, effects from stimulating the two sites in humans differed. We calculated the passive current spread as above: resulting in a 3 mm radius around the stimulation contact pairs, which are separated by 5-10 mm. The analytical group contrast of Site 1 (medHG) vs Site 2 (latHG+PT) showed stronger vlPFC activity from stimulating Site 1 (Fig. 3; Table 2). Site 1 stimulation also resulted in stronger activity in more posterior temporal and parietal areas (e.g., supramarginal gyrus). Site 2 stimulation resulted in stronger activity in anterior temporal areas (Table 2).

**Figure 3.**
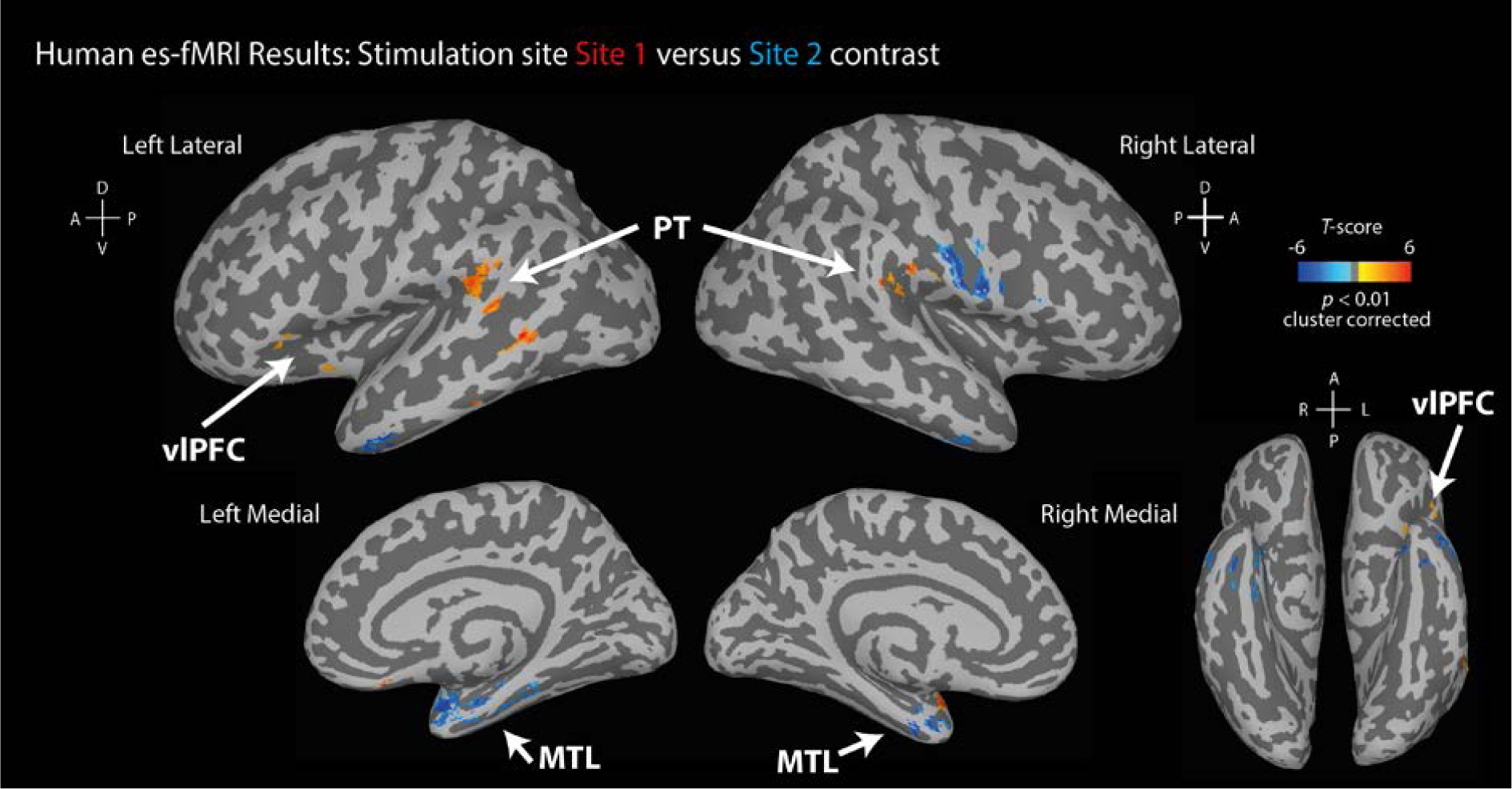
Human results contrasting Site 1 versus Site 2 esfMRI effects. Same format as in Fig. 2C-D, showing statistically significant (*p* < 0.01 cluster-corrected) effects where either Site 1 (red color map) or Site 2 (blue color map) were stronger. The corresponding contrast in monkeys yielded no cluster-corrected differences.

### Differential macaque esfMRI effects in vlPFC and MTL subregions

We assessed whether macaque auditory cortex stimulation differentially activates anatomically defined areas 44, 45 and the FOP in the vlPFC (Fig. 4A-B). Region of interest (ROI) effects were tested with a mixed-design analysis of variance (ANOVA): between-subjects factor of Monkeys, within-subjects factors of Activated hemisphere (left or right), ROI (area 44, 45 or FOP) and Stimulation site as covariate (Site 1 or 2). Model assumptions for this and all subsequent reported analyses were met or corrected as indicated. Planned post-hoc tests were used to identify differential effects across the ROIs.

**Figure 4.**
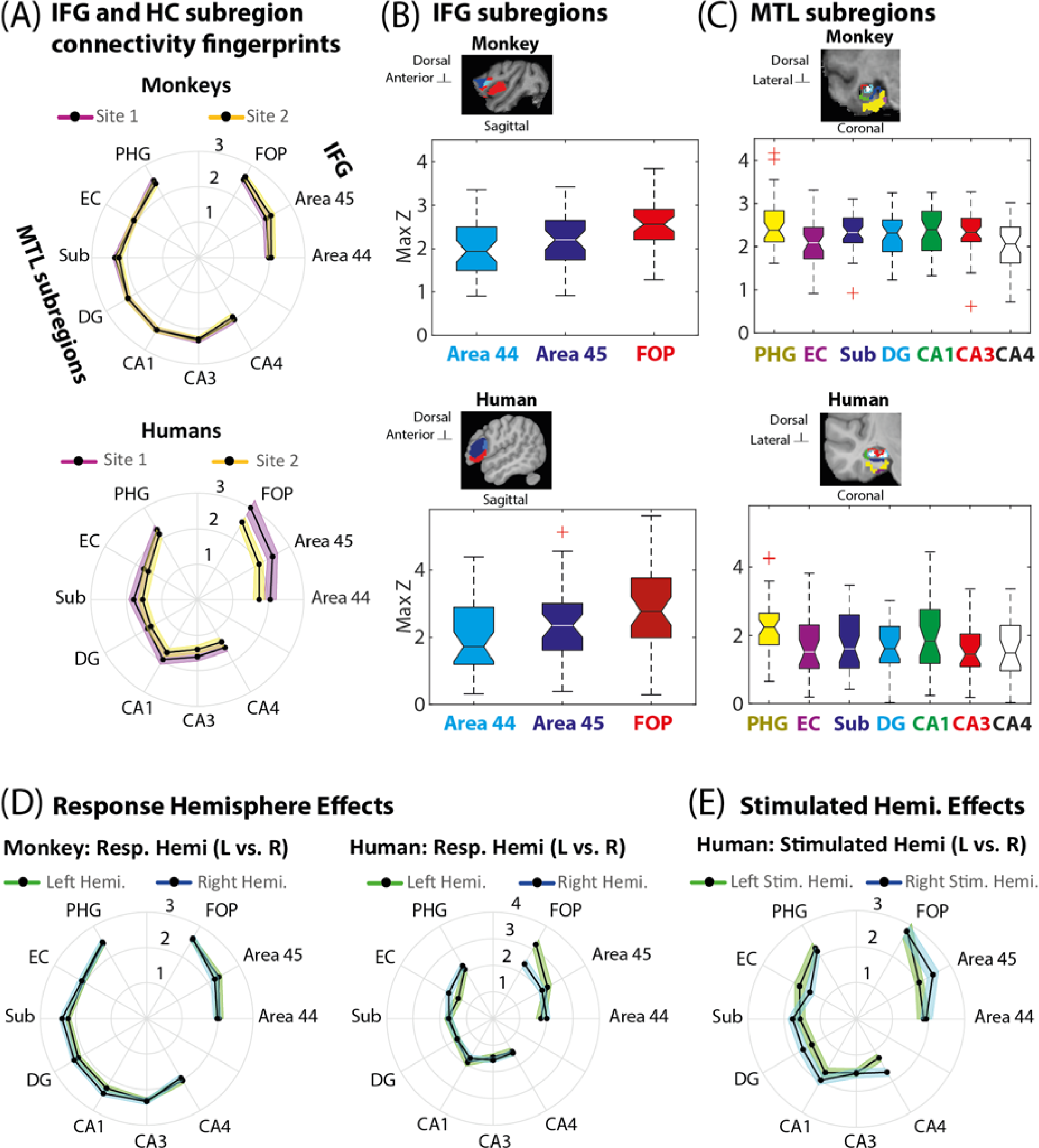
Human and macaque vlPFC and MTL connectivity profiles. (A) vlPFC and MTL subregion esfMRI effects displayed as polar plots. Shown are the across scanning runs peak *z*-values and variability (+/-SEM, standard error of the mean). Top plot in (A) shows monkey results, bottom human results. (B-C) whisker plots of vlPFC (B) and MTL (C) esfMRI activity responses (across scanning runs, peak z-value; central mark identifies the median, edges of box are 25^th^ and 75^th^ percentiles, whiskers extend to extreme ends of data not including outliers in red crosses; non-overlapping notches are significantly different at *p* < 0.05). Also shown are sagittal and coronal slices in each species with the anatomically localized ROIs used for the analysis. (D) effects by response hemisphere (monkeys left, humans right). (E) human effects by stimulated hemisphere; only right hemisphere was stimulated in the monkeys.

The overall magnitude of the fMRI signal in the vlPFC ROIs was stronger in macaque 1 (M1; significant Monkey factor: *F*_1,34_= 7.34, *p* = 0.010), but there were no interactions with other factors, suggesting a similar pattern of esfMRI effects across the monkeys. The two Sites of stimulation did not differentially affect the vlPFC fMRI response, consistent with the whole-brain results. The macaque vlPFC ROI effects were also statistically indistinguishable across the two hemispheres (no significant Hemisphere effect). The ROIs showed a trend towards a differential fMRI response (*F*_2,33_ = 3.04, *p* = 0.054), and the planned post-hoc comparisons identified stronger esfMRI activity in the FOP over areas 45 and 44 in that order (all *p* < 0.002, Bonferroni corrected; Fig. 4A-B).

Monkey MTL subregions showed differential esfMRI activity during auditory cortex stimulation (Fig. 4C). A mixed-design ANOVA including the seven anatomically delineated MTL subregions showed that esfMRI activity in these ROIs did not differ between monkeys. Effects also did not differ across the two stimulation sites, or across the two hemispheres. The MTL ROIs differed nonlinearly in their fMRI activity response (cubic effect: *F*_1,34_ = 4.27, *p* = 0.028; Fig. 4A, C). The planned post-hoc comparisons showed stronger esfMRI activity in parahippocampal gyrus (PHG) than entorhinal cortex (EC) and CA4 (*p* < 0.003, Bonferroni corrected). Subiculum (Sub), dentate gyrus (DG), CA1 and CA3 also had stronger activity than CA4 (all *p* < 0.015, Bonferroni corrected).

### Differential human auditory cortex esfMRI effects in vlPFC and MTL

We studied the human vlPFC and MTL effects in anatomically defined ROIs. Effects were tested with mixed-design ANOVA (between-subjects factor: Human subject; within-subject factors: ROI subregion, Activated hemisphere, left or right; Site of stimulation and Stimulated hemisphere, left or right, as covariates). The vast majority of subjects were left hemisphere language dominant (Table 3).

**Table 3.** Human patient demographics

| Patient ID | Age | Sex | Handedness | Language dominance (Wada test) | esfMRI | esT | Speech |
| --- | --- | --- | --- | --- | --- | --- | --- |
| 292 | 50 | F | L | L | Yes (Y) |  |  |
| 302 | 47 | F | R | ** | Y |  |  |
| 307 | 30 | M | R | L | Y |  |  |
| 314 | 30 | F | R | L | Y |  |  |
| 316 | 31 | F | R | ** | Y |  |  |
| 320 | 50 | F | R | ** | Y |  |  |
| 330 | 43 | M | L | ** | Y |  |  |
| 331 | 35 | M | R | ** | Y |  |  |
| 334 | 39 | M | L | L | Y |  |  |
| 335 | 31 | M | R | L | Y |  |  |
| 339 | 45 | M | R | ** | Y |  |  |
| 352 | 31 | M | Mixed | ** | Y |  |  |
| 357 | 36 | M | R | L | Y |  | Y |
| 369 | 30 | M | R | L | Y | Y | Y |
| 372 | 34 | M | R | L | Y | Y | Y |
| 376 | 48 | F | R | L | Y | Y | Y |
| 384 | 37 | M | R | L |  | Y |  |
| 394 | 23 | M | L | L |  | Y | Y |
| 395 | 13 | M | R | ** | Y |  |  |
| 399 | 22 | F | R | L | Y | Y | Y |
| 400 | 59 | M | L | L | Y |  |  |
| 403 | 56 | F | R | L | Y |  |  |
| 405 | 19 | M | R | L | Y | Y | Y |
| 413 | 22 | M | L | R | Y | Y |  |
| 418 | 25 | F | R | L |  | Y |  |
| 423 | 49 | M | R | L |  | Y | Y |
| 427 | 17 | M | Mixed | ** |  | Y |  |
| 429 | 32 | F | R | ** |  | Y | Y |
| 457 | 18 | M | R | ** |  |  | Y |
| 460 | 52 | M | R | L (fMRI) |  | Y |  |

\*\* (not done)

For vlPFC, we observed differential activity of areas 44, 45 and FOP (*F*_1,21_ = 5.26, *p* = 0.032; Fig. 4B) in a similar pattern as in the monkeys. The post-hoc tests showed a stronger FOP response than in areas 45 and 44 (*p* < 0.003, Bonferroni corrected) with a trend for stronger area 45 than area 44 activity (*p* = 0.070). Site of stimulation (Site 1 vs Site 2) effects on vlPFC were a trend (*F*_1,21_ = 3.14, *p* = 0.090), with a borderline weaker Site 2 (latHG+PT) esfMRI effect on the vlPFC subregions (Fig. 4A). Hemispheric differences for certain ROIs were evident: visualized in Fig. 4D as a stronger FOP response on the left, with the right hemisphere showing similar activity across the three ROIs. Activated hemisphere effects did not interact with vlPFC ROI but did with Site of stimulation (Site 1 vs Site 2 by hemisphere interaction: *F*_1,21_ = 7.23, *p* = 0.014) and Stimulated hemisphere (Stimulated hemisphere by Activated hemisphere interaction: *F*_1,21_ = 4.68, *p* = 0.042; Fig. 4E).

For the human MTL ROI effects, the ANOVA did not show significant differential activation between the ROIs, although the albeit weaker pattern was similar to that seen in monkeys with the following post-hoc comparisons: The PHG had a stronger esfMRI response than CA4 (*p* = 0.016, Bonferroni corrected) and DG (*p* = 0.026, corrected), and the subiculum response was stronger than CA4 (*p* = 0.034, corrected). MTL ROI effects showed a trend for Activated hemisphere (*F*_1,10_ = 3.35, *p* = 0.097), seen in Fig. 4D as a slight right hemisphere bias particularly for entorhinal cortex (Fig. 4D-E). No other significant effects or interactions were observed.

### Cross-species auditory cortex esfMRI comparisons

We conducted cross-species esfMRI comparisons of the vlPFC and MTL ROI responses (Fig. 4 shows the anatomically-delineated ROIs in both species). Statistical testing included Species as a between-subjects factor.

For the vlPFC cross-species comparison, both monkey and human esfMRI results showed stronger FOP responses than area 45 (ROI effect: *F*_1,57_ = 9.68, *p* = 0.003). There was no significant species difference in the vlPFC ROI effects (*p* = 0.139), showing statistically indistinguishable vlPFC ROI response patterns across the species (Fig. 4A-B). An effect was found for Activated hemisphere (left stronger than right, *F*_1,57_ = 4.21, *p* = 0.045) but did not significantly interact with Species. There were higher order interactions with species, including hemispheric differences (vlPFC ROIs by Species: *F*_1,57_ = 4.97, *p* = 0.010; ROIs by Activated hemisphere by Species: *F*_2,57_ = 7.11, *p* = 0.002).

For the MTL, the overall esfMRI activity level differed across the Species (Fig. 4D, *F*_2,46_ = 10.55, *p* < 0.001). The ROIs did not differ in their fMRI response pattern but interacted with Species (Greenhouse Geisser corrected: *F*_7.63,175.52_ = 3.05, *p* = 0.004, ε = 0.636), and stronger hemispheric lateralization was seen in humans (Species by Activated hemisphere interaction; Greenhouse Geisser corrected: *F*_8.70,200.16_ = 2.175, *p* = 0.027, ε = 0.725). For instance, entorhinal cortex shows greater right hemisphere activation (Fig. 4D), and this region was relatively more activated when the left auditory cortex was stimulated (Fig. 4E). By contrast, the monkey effects were statistically indistinguishable across the two hemispheres. No other effects or higher-order interactions were significant.

### Human electrical stimulation tractography (esT)

In the human patients, clinical electrode coverage could include vlPFC, auditory cortex and hippocampus. We studied neurophysiological connectivity with esT between auditory cortex, vlPFC and hippocampus (Figs. 5A, F-H show the stimulation sites and recording contacts).

Auditory cortex Site 1 (MedHG) and Site 2 (latHG+PT) stimulation induced neurophysiological potentials in vlPFC (Fig. 5B-C) as soon as they could be measured after the 0-3 ms electrical stimulation artifact (Fig. 5D; Suppl. Movies 1-6). We assessed the polarity and latency of stimulation-induced responses using spatial Laplacian correction to reduce volume conduction effects and spurious cross-channel correlated responses. Response waveforms in vlPFC showed positive and negative components at different latencies (Fig. E-H). Our labelling convention follows prior reports^16,28^, identifying positivities (*P*) and negativities (*N*) by latency (early: *a*; later: *b*).

**Figure 5.**
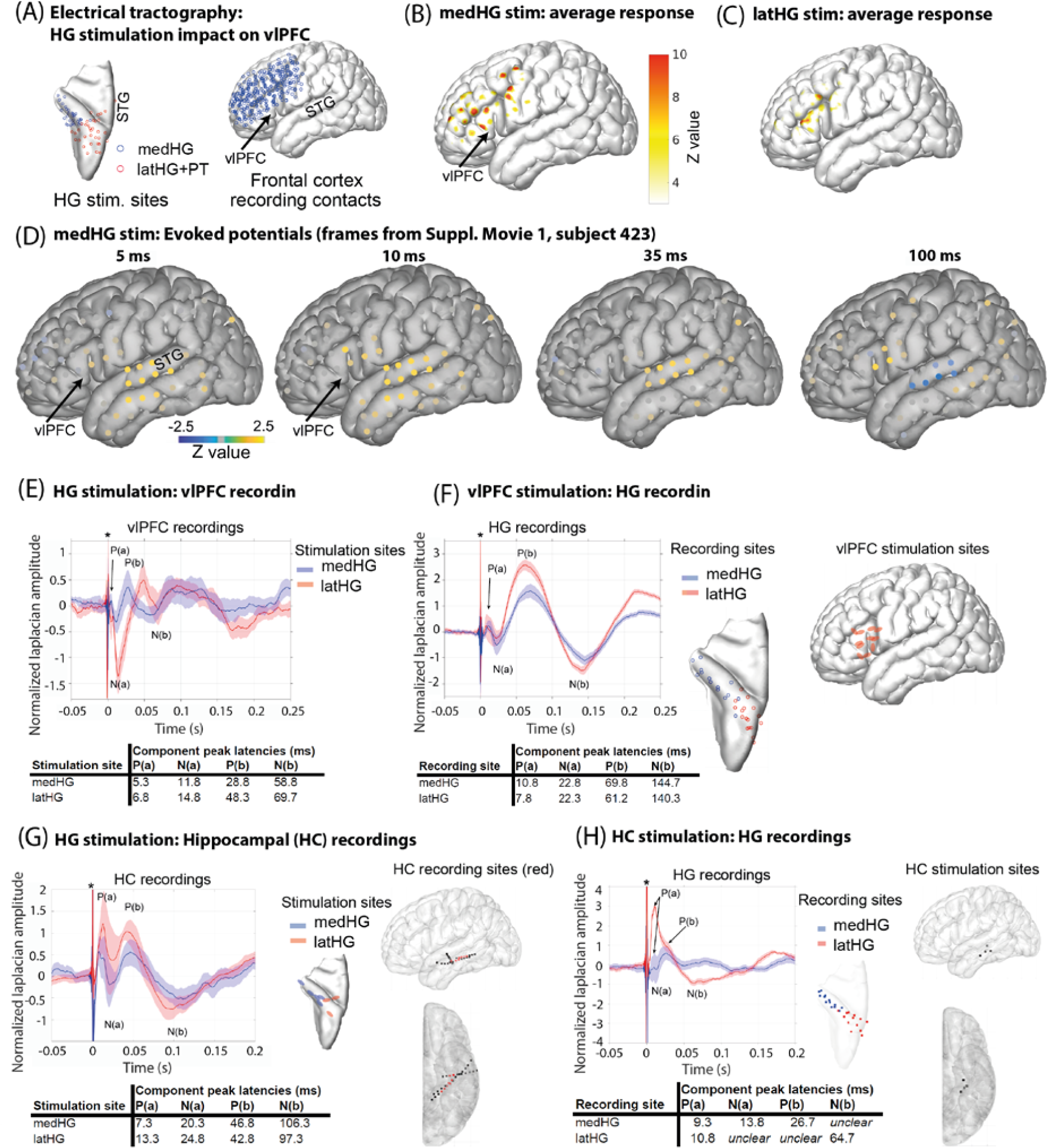
Human electrical tractography. (A) Human HG stimulation sites and vlPFC recording electrode locations. (B-C) average gamma band evoked response from the recording contacts shown by medHG (B) or latHG+PT (C) stimulation. (D) frames at 5, 10, 35 and 100 ms post-electrical pulse stimulation from Supplementary Movie 1 (subject 423) during medHG stimulation. (E) average neurophysiological evoked potentials in vlPFC from stimulating the HG sites, showing the peak latency (ms) for each component. (F) vlPFC stimulation and recording in HG; same format as in (E); location of stimulation and recording contacts shown on right. (G) stimulation of HG during recordings in hippocampus (HC). (H) stimulation of HC during recordings in HG.

Auditory cortex stimulation induced an early vlPFC positivity (*P(a)*: average latency: 5.3 and 6.8 ms, respectively from stimulating medHG or latHG+PT) followed by an early negativity, *N(a)*, and later positivity and negativity (Fig. 5E). Notably, the early positivity is as early as reported from stimulating medial HG and recording in postero-lateral STG^28^ (Suppl. Fig. S8 shows replication).

Effects from stimulating vlPFC while recording in HG indicate that connectivity appears to be bi-directional, evident as the presence of both sets of positivities and negativities: compare Fig. 5F with stimulation in the opposite direction in Fig. 5E. However, the waveforms do differ particularly in the latencies of some of the components which are earlier for the HG to vlPFC direction. An ANOVA (within-subjects factor: Potential latency across recordings, between-subjects factors: Stimulation sites and Subjects) substantiated this, showing a significant interaction of Stimulation site with Potential latency (Greenhouse Geisser corrected: *F*_1.467,77.742_ = 45.641, *p* < 0.001, ε = 0.489). No other effects or interactions were found.

Next we studied esT effects between hippocampus and auditory cortex. The waveforms recorded in the hippocampus after stimulation of HG, or vice versa, were also distinctly different in shape, with later components much more variable in latency depending on the direction of stimulation (compare *N(b)* latencies in Fig. 5G-H). There was a significant interaction of Stimulation site with Potential latency (Greenhouse Geisser corrected: *F*_1.407,45.034_ = 9.399, *p* < 0.001, ε = 0.469). No other significant effects or interactions were found.

Testing only the earliest *P(a)* component (excluding the others), showed it to be significantly earlier when stimulating HG and recording in vlPFC than any of the other combinations of directions or stimulation/recording sites (all comparisons relative to stimulating HG and recording in vlPFC, *p* < 0.001, Bonferroni-corrected). Notably, this early latency response in vlPFC resulting from stimulating HG appears as early as reported from potentials recorded in the posterior STG when stimulating HG^28^ or when recording in vlPFC and stimulating STG^16^.

### Human vlPFC speech responses and directional connectivity with auditory cortex

Lastly, we studied whether speech sounds induce neurophysiological responses in human vlPFC or MTL and the directionality of neurophysiological interactions using state-space Conditional Granger Causality (CGC). Expectedly, auditory (HG and STG) sites showed strong broad-band speech-driven responses (Fig. 6A), including power decreases in low frequencies after speech onset^29^. Individual contacts showed significant speech responses in PHG and hippocampus, although responses in these areas were weak in the group average results. By comparison, vlPFC responses to speech sounds was much more substantial in individual and group results (Fig. 6A), evident as increases in lower frequency (theta) power with suppression in alpha and beta bands.

**Figure 6.**
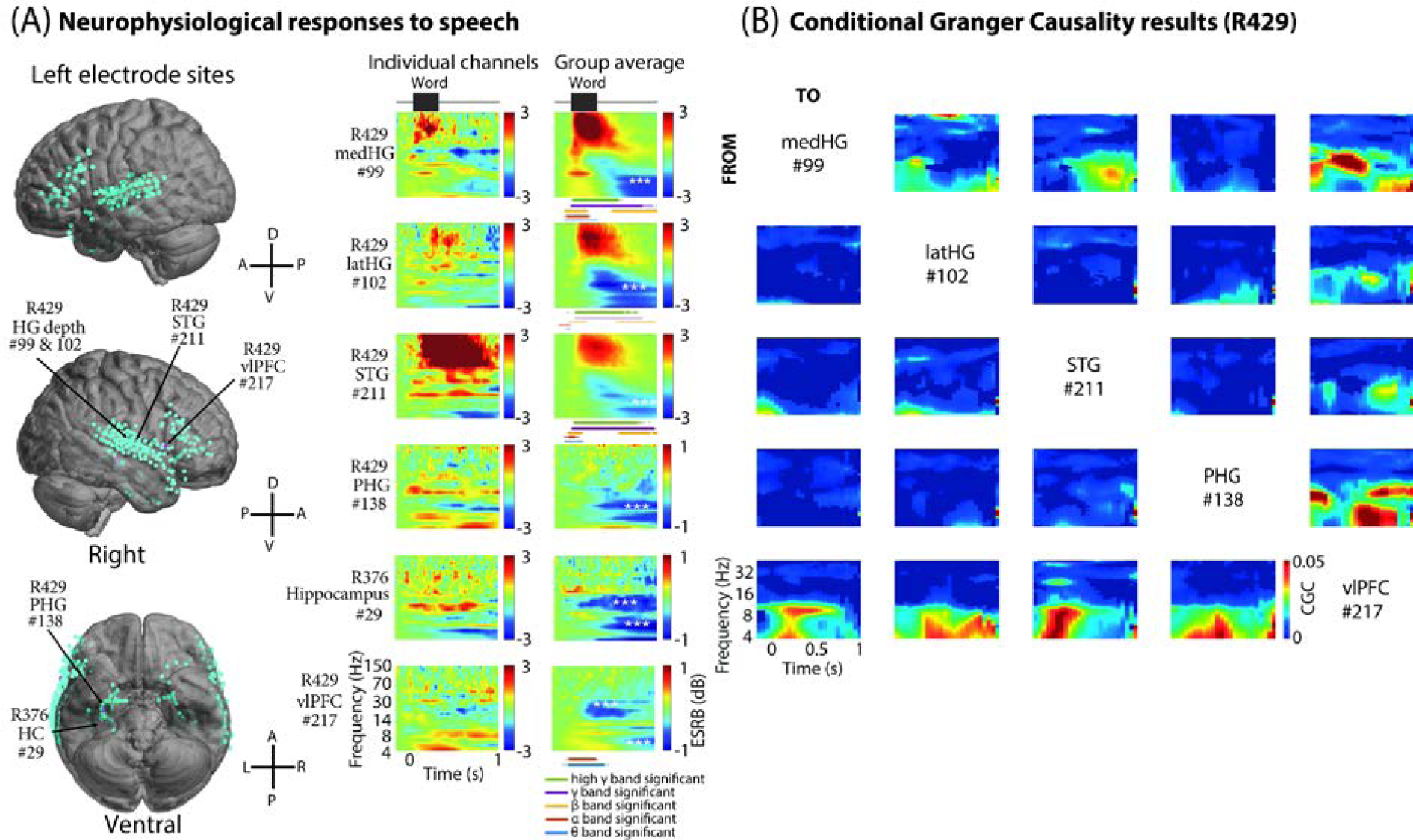
Human vlPFC responses to speech and conditional Granger causality interconnectivity with auditory cortex. (A) left, electrode locations across all participants (*N* = 8) projected on the left and right lateral and ventral view a template brain across all the participants. Right, time-frequency resolved responses to speech sounds (common words). Shown are single subject individual channels (left column) and group average (right column) within Heschl’s gyrus (HG), superior temporal gyrus (STG), parahippocampal gyrus (PHG), hippocampus (HC) and vlPFC. Subject 429 did not have hippocampal coverage, therefore the responses from another subject (376) are shown for this region. Below the group results we identify significant responses subdivided by frequency bands (thin > 2 SD; thick > 4 SD relative to the pre-word baseline variability). White *** symbols inset in the group plots identify significant suppression. (B) state-space Conditional Granger Causality (CGC) results showing directional neurophysiological interactions during speech sound presentation. Directions of influence are shown from regions of interest (rows) to recipient regions (columns) active during speech presentation. Subthreshold (not significant) regions of time-frequency CGC were set to 0 and are masked in dark blue. Note the strong dynamic directional influences particularly between vlPFC and auditory sites such as HG.

Directional frequency-resolved CGC analyses were conducted between pairs of contacts with the results conditioned on non-specific effects across all contacts during the neurophysiological responses to speech (subject 429, Fig. 6B). Expected HG and STG directional interactions were observed (Fig. 6B). In this subject there were no hippocampal contacts, but the available PHG contact only showed significant directional interconnectivity with vlPFC; PHG connectivity with auditory cortex was weak. By comparison, the vlPFC showed strong bi-directional interactions with the auditory sites (Fig. 6B).

## DISCUSSION

In monkeys, neuronal tractography has shown both direct projections from non-primary auditory cortex to vocalization sensitive vlPFC neurons^14,15,27^ and indirect projections from auditory cortex to the hippocampus via parahippocampal cortex^20,21,30,31^. In humans, because of language and declarative memory the corresponding pathways are expected to have specialized in our species. Amidst comparative insights on differentiation involving language pathways^7-9,32^, there is surprisingly little evidence for a privileged auditory to vlPFC pathway in humans^7,16,17,33^ as in macaques^14,15,27^. Also, human resting-state data is suggestive of indirect auditory to hippocampal functional connectivity^23,34^, as in other species, with the only evidence for direct projections between hippocampus proper and hierarchically earlier auditory fields having been obtained in rodents^22^.

We harnessed a new approach for similarly assessing effective connectivity in humans and monkeys, using combined electrical stimulation and functional MRI (esfMRI)^26,35^. We found comparable fronto-temporal effective connectivity in both species from stimulating auditory cortex in several vlPFC and MTL subregions. More subtle differences in human hemispheric lateralization were seen, relative to more bilaterally symmetrical effects in monkeys. In humans, we also obtained evidence for rapid electrically-induced neurophysiological responses in vlPFC from stimulating auditory cortex, with shorter latency in the first measurable potential than that seen in the opposite direction (stimulating vlPFC and recording in the auditory HG), or for interactions between HG and hippocampus. Also, we identified a speech responsive region in human vlPFC with directional effective connectivity with auditory cortex. The findings show human auditory pathways that appear to be as direct to vlPFC and indirect to the hippocampus (via parahippocampal cortex) as in nonhuman primates, illuminating the primate origins of fronto-temporal pathways for cognition.

### Common auditory esfMRI fronto-temporal effective connectivity

The comparison of esfMRI effects in humans and monkeys supports the notion of fronto-temporal networks including auditory cortex, vlPFC and MTL being based on a largely evolutionarily conserved effective connectivity signature. In both species, we observed stronger frontal operculum (FOP) responses than in vlPFC areas 44 and 45, with some hemispheric difference in humans, evident as a more balanced response across these regions in the human right hemisphere. Also in both species, auditory cortex stimulation induced stronger activation in PHG than other MTL subregions.

The strong FOP effect is interesting because this region ventral to vlPFC areas 44 and 45 has been implicated in initial syntactic processes in humans, based on its involvement in processing local relationships such as those between adjacent words in a sentence^9^. Comparative human and monkey fMRI using a sequence learning task has shown that the FOP, in particular, similarly responds in these species to adjacent sequencing relationships^36^. Also, the FOP is part of a fronto-temporal saliency in attention or task control network that appears to be evolutionarily conserved in primates^37^.

The comparably stronger esfMRI effect in monkeys and humans from stimulating auditory cortex seen in the PHG, which was stronger than certain hippocampal subregions, suggests an equally indirect auditory cortical projection to the hippocampal memory circuit across these species. This observation is not inconsistent with impressions from human resting-state connectivity, including ultra-high-resolution data^33,34^. The current findings challenge the notion that monkeys have a less direct auditory effective interconnectivity with the MTL memory system than humans. Our language abilities allows us to name, conceptualize and thus better remember sounds^24^, but, as our observations suggest, possibly with a largely evolutionarily conserved MTL system.

The mode of action of esfMRI remains poorly understood. A prior monkey esfMRI study showed that electrical stimulation of the visual thalamus elicits an fMRI response in primary visual cortex (V1) with inhibitory neurons reducing signal propagation to other areas^38^. However, some propagation to multi-synaptically connected sites appears to be evident in the form of graded fMRI responses. Namely, our pattern of graded MTL ROI effects are generally consistent with known parahippocampal and hippocampal subregion multi-synaptic connectivity^39,40^.

### Cross-species differences in esfMRI effects

Evidence for human differentiation was clearest in the form of hemispheric lateralization. By contrast, the monkey esfMRI effects were largely bilateral (even though it was only possible to stimulate the right hemisphere in the monkeys). The human results showed hemispheric lateralization effects for vlPFC and MTL subregions, almost independently of which hemisphere was stimulated (Fig. 4D).

There were also differences in effects between humans and monkeys by auditory cortical site of stimulation. We initially stimulated auditory ‘core’ and ‘belt’ areas in the monkeys aiming to identify distinctly different esfMRI effects, given that belt neurons are known to project to prefrontal cortex^41^. However, the monkey results were statistically indistinguishable from stimulation of these two adjacent auditory sites, potentially because passive current spread was estimated to be ∼1 mm radius around the stimulating electrode (Results). The human results were distinctly different between the two stimulation sites, one of which was identified by its auditory click following response^42^. Passive current spread in the human results is also worth considering, with ∼2.5 mm radius around the stimulating contacts, indicating that some of the human medHG effects can involve stimulation of adjacent areas.

### Considerations in relation to white matter pathways and monkey anterograde tractography

The effective connectivity approach we used is agnostic to the white matter tracts that interconnect auditory cortex with the fronto-temporal sites. Human diffusion MRI and monkey anterograde tractography indicate that the FOP is interconnected with auditory cortex via the ventral extreme-capsule or uncinate fasciculus pathway^43^. Human area 44 is interconnected with areas caudal to auditory cortex via the dorsal arcuate fasciculus pathway^13^. A recent comparative study showed a homolog of this dorsal auditory pathway in macaques^8^ projecting from caudo-medial regions near to those that we electrically stimulated, which elicited vlPFC activity.

Is it unexpected that our esfMRI results from stimulating posterior auditory cortical sites show effects in vlPFC? Namely, monkey anterograde tractrography results from caudal auditory belt areas (like area CL) show denser *dorso-lateral* prefrontal cortex axonal bouton labelling^15,27^. Although mesoscopic effects such as those measured by esfMRI lack the specificity of single neuron tractography, our results do recapitulate key patterns shown with macaque anterograde tractography, such as both anterior and posterior auditory belt neuronal tracer injections labelling axonal boutons in vlPFC^15^. Also, in relation to a prior macaque esfMRI study, our results are more similar to their effects from stimulating posterior lateral belt (area ML) than those from stimulating the anterior lateral belt (RTL)^44^. In that study there was one auditory cortical field separation between the two stimulated sites. In our study, there was no such buffer region between the two stimulated sites, which may be why our effects from stimulating the two sites were statistically indistinguishable. Another consideration is the inherent differences in how the monkeys and humans were studied (e.g., different scanners, electrodes, parameters). However, amidst all such differences, it is remarkable how similar the esfMRI effects in vlPFC and MTL were across the species.

### Rapid human auditory cortical to vlPFC electrical tractography

The study provides evidence for a rapid electrical-tractography response in vlPFC following HG stimulation. An early vlPFC positive potential occurred as soon as we could measure after the stimulation artifact, peaking ∼5-7 ms after HG stimulation. Auditory cortex and the vlPFC are separated by ∼10 cm via the arcuate fasciculus. The latency of this early positivity was significantly longer in the opposite direction (stimulating vlPFC and recording in HG) and between HG and the hippocampus in either direction (Fig. 5E-H). The estimated conduction velocity, based on axonal diameter and conduction speeds^45,46^ including within fronto-temporal pathways^47^, is 10-30 m/s. Thus, the expected latency of the first potentials from auditory cortex arriving in vlPFC is between ∼3-10 ms. Our HG to vlPFC latency results are remarkably similar in latency to the reported ∼6 ms peak of the initial positivity recorded in auditory posterior STG sites after medial HG stimulation^28^; the two sites are separated by 1-2 cm. The authors in that study could not exclude the possibility of a monosynaptic connection between medial HG and the STG, because, surprisingly, stimulation of a lateral HG site expected to be an intermediary elicited weaker recorded potentials in the STG. In another study, stimulation of the posterior STG while recording in vlPFC resulted in an early negativity (the study focused on negative potentials) with an average vlPFC response latency of ∼13.5 ms^16^, which is similar to our first negativity latency in vlPFC from stimulating HG (*N(a)*; Fig. 5E). Even longer first negativity latencies in vlPFC can be seen from stimulating various temporal lobe sites^17^.

Thereby, our electrical tractography effects from HG to vlPFC are notably similar to those reported from recording in vlPFC after stimulating STG^16^. The results raise the possibility that the HG to vlPFC connection is as rapid as the one from posterior STG to vlPFC. Although we cannot exclude multi-synaptic effects, it is possible that if medial or lateral HG sites require additional synaptic connections in the STG, that the vlPFC early positivity (or negativity) could peak later. Moreover, although HG and vlPFC effective connectivity appeared to be bidirectional, there were key differences in the shape of waveforms in both directions and the early positivity was earlier when stimulating HG and recording in vlPFC than in the opposite direction. The early positive potential between the HG and hippocampus in either direction was also later than the one observed between HG and vlPFC. Our latencies between HG and the hippocampus are however within the range of those reported from recording in various temporal lobe sites after entorhinal cortex stimulation^48^. The results provide support for privileged HG to vlPFC effective connectivity that differs in the opposite direction and from results involving the hippocampus.

### Human speech responsive vlPFC region with directional auditory cortex effective connectivity

We observed considerable speech sound driven neurophysiolocal responses in vlPFC and show conditional Granger causality results on bidirectional effective connectivity between HG and vlPFC. Speech responses in vlPFC are not unexpected, typically being evident during active speech recognition or difficult listening conditions^49^. Spoken speech or reading can also elicit responses from the human hippocampus^50^ (see Fig. 6). The conditional Granger causality results showing interconnectivity between vlPFC and HG during speech processing is further evidence for a privileged auditory to vlPFC pathway in the human brain.

In summary, the findings demonstrate largely comparable effective connectivity signatures between human and macaque auditory cortex, vlPFC and MTL. The auditory system as the model sensory system under study here is expected to show human-specific specialization for speech, language and declarative memory. However, even these results involving auditory cortex show considerable correspondence in fronto-temporal effective connectivity across these species. Future studies in other species or sensory modalities could further support or refute our observations.

## ACKNOWLEDGEMENTS

Supported by Wellcome Trust (CIP: WT092606AIA; TDG: WT091681MA), Biotechnology and Biological Sciences Research Council UK (CIP: BB/J009849/1), National Institutes of Health USA (MAH: R01-DC04290; JDWG: R01-DC015260; RA and MAH: U01-NS103780) and European Research Council (CIP: ERC CoG, MECHIDENT). We thank Lauren Dean and Jennifer Nacef for assistance with the macaque data collection, and Haiming Chen, Phil Gander and Beau Snoad for assistance with the human data collection.

## AUTHOR CONTRIBUTIONS

Conceptualization: CIP, MAH, TDG. Investigation: FR, HO, FB, AER, KVN, MS, YK, HK, CIP. Formal Analysis: FR, HO, ZK, RJ, CIP. Resources: FB, AB, CKK, KVN, YK, BJD, HK, RA, JDWG, TDG, MAH, CIP. Writing – Original Draft: FR, HO, CIP. Writing – Review & Editing: FR, HO, FB, AB, ZK, RJ, KVN, YK, BJD, RA, JDWG, TDG, MAH, CIP. Supervision: MAH & CIP. Project administration: HO, FR, FB, AER, KVN, MAH, CIP. Funding acquisition: RA, JDWG, TDG, MAH, CIP.

## ONLINE METHODS

### Macaque procedures

All the nonhuman animal work and procedures were approved by the university Animal Welfare and Ethical Review Body and UK Home Office. The work complies with the revised UK Animal Scientific Procedures Act (1986), the US National Institutes of Health Guidelines for the Care and Use of Animals for Experimental Procedures and with the European Directive on the protection of animals used in research (2010/63/EU). We support the principles on reporting animal research stated in the consortium on Animal Research Reporting of In Vivo Experiments (ARRIVE). All persons involved in animal handling and procedures were certified and the work was strictly regulated by the UK Home Office.

Two male rhesus macaques (*macaca mulatta*) from a group-housed colony were scanned awake with combined electrical stimulation and fMRI (M1 and M2, 10 and 12 years old, weighing 12.2 and 12.6 kg, respectively). The pen sizes in the colony range from 130 x 240 cm to 215 x 240 cm. All are 230 cm high, and hatches between neighboring cages are used to increase the space available to the animals. Given the ethical sensitivities involved in studying nonhuman primates and the 3Rs principles (one of which is on the Reduction of animal numbers), our work requires using the fewest monkeys possible. A sample size of two is common in neuroscience work with macaques provided that results are robust with each individual and that the effects generalize beyond one animal. Given that our results from several hundred trials and many scanning runs with each animal are statistically robust and consistent in the overall pattern of effects between the animals (i.e., no significant interactions of monkeys as a factor with the reported patterns of esfMRI results) there was little ethical justification to test additional monkeys.

### Macaque electrical stimulation procedure

The electrical stimulation procedure used in this work is based on methodology developed in prior macaques esfMRI studies^25,38,44,51^. Here the procedure was conducted on awake monkeys in absence of alterations due to anesthetics^52,53^ to allow more direct comparison with awake human esfMRI data.

Prior to the experiments, an MRI-compatible PEEK head post and chamber were implanted stereotaxically during an aseptic procedure under general anesthesia. The chamber was positioned over the right hemisphere to provide access to posterior auditory cortex, including fMRI tonotopically localized fields A1 and the posterior area adjoining the border between belt fields CL and CM (Suppl. Fig. 1).

During each scanning session, a custom-made PEEK microdrive was used to advance the stimulating electrode to auditory cortex. Electrical stimulation was induced through platinum/iridium microelectrodes coated in parylene-C (Microprobes, Gaithersburg, Germany), with a typical impedance of 100-200 kΩ (electrodes with impedance below 75 kΩ were not used). The stimulating electrode location in auditory cortex was confirmed via MRI structural scans and sound-evoked neurophysiological responses as in previous studies^44^, using a TDT system running Synapse (Tucker-Davis Technologies, Alachua, FL). The experiment was controlled by Matlab software (MathWorks, Natick, Massachusetts, US) running the Psychophysics Toolbox, interacting with the hardware connected via a LabJack interface device (www.labjack.com). The experimental computer triggered each EPI scan.

The target areas were electrically stimulated using a World Precision Instruments (DS8000) waveform generator with an electrical stimulus isolator unit (DLS 100). The system delivered a constant-current, charge-balanced electrical pulse of either 0.5 or 1 mA (fixed within a stimulation session) at a repetition rate of 200 Hz (within 4 pulse periods; Fig. 1A). Each biphasic-square-wave pulse had 0.2 ms duration. The stimulation and MRI scanning timing paradigm are shown in Fig. 1B. Electrical stimulation was randomly induced in 50-70% of the scanning trials with the others containing no stimulation, as a baseline point of reference.

### Macaque fMRI

Functional MRI measured the blood oxygen level dependent (BOLD) signal, using an actively shielded vertical primate-dedicated 4.7 Tesla MRI scanner (Bruker BioSpin, Ettlingen, Germany). The monkeys had been slowly acclimated over several months with reinforcement training to work in the primate scanning chair and with the required periods of head immobilization during fMRI^54^. They were scanned awake under head immobilisation while conducting a visual spot fixation task^36^.

A custom 4-channel surface receiver coil array and a saddle transmitter coil were used for MRI acquisition (WK Scientific, California USA). Functional data were obtained using a gradient-recalled echo planar imaging (GE-EPI) sequence, with the following parameters: echo time (TE) = 20 ms; volume acquisition time (TR) = 2000 ms; flip angle = 90 degrees; 32 slices, in plane with a field of view (FOV) of 10.7 × 10.7 cm^2^, on a grid of 88 × 88 voxels. Voxel resolution was 1.2 × 1.2 × 1.3 mm^3^ covering much of the brain. A sparse fMRI acquisition paradigm was used to minimize the interference caused by the scanner noise on the auditory cortex response^55,56^; a single T2* weighted MRI volume was acquired after the electrical stimulation or no-stimulation periods for each trial. The onset of the volume lagged by ∼4 s to account for the lag in the hemodynamic response^57^. The number of trials obtained per scanning run for each animal were 60 for M1 and 120 for M2 and the numbers of scanning runs analyzed were 20 (9 in M1; 11 in M2) for Site 1 stimulation and 17 (8 in M1; 9 in M2) for Site 2, see Fig. 1.

Anatomical images, including those for helping to visualize the electrode position, were acquired at the beginning and end of each experimental testing session using magnetization-prepared rapid gradient-echo (MP-RAGE) sequences. Typical sequence parameters were: TE = 20 ms; inversion time, TI = 750 ms; TR = 2000 ms; 50 slices with in-plane field of view: 10.7 × 10.7 cm^2^ on a grid of 140 x 140 voxels. The voxel resolution was 0.75 × 0.75 × 0.75 mm^3^.

### Macaque fMRI data analysis

We performed General Linear Analysis using FEAT in FSL^58,59^. We contrasted the BOLD responses during stimulation and non-stimulation trials. Each scanning volume in this sparse imaging paradigm was assigned to either the stimulation or no-stimulation conditions. Brain extraction used the BET function in FSL, and the fMRI image series were motion corrected within a given testing session using FLIRT (typically 9 or 12 degrees of freedom affine transformation with the normalized mutual information option). The motion parameters were used as regressors of no interest in the fMRI analyses. The data were smoothed with a Gaussian kernel 2 mm Full Width Half Maximum (FWHM). For registration of the fMRI time series, the T2-weighted scans were registered to the animal’s session full-brain mean functional scan. This scan was then registered to the animal’s session anatomical and then to its high-resolution anatomical scan. Finally these scans were registered to a macaque template brain^60^, which is in register to a macaque atlas in stereotactic coordinates^61^.

*General Linear Modelling:* We first preprocessed each scanning run using first-level analyses. Individual scanning runs with little evidence of the expected electrically induced activity (at a liberal uncorrected threshold *Z* > 2.3) in auditory areas around the electrode or in auditory cortex in the opposite hemisphere were not analyzed further. Group higher-level analyses were conducted combining all of the viable scanning runs grouped by stimulation site (Site 1 or 2). Higher-level analyses were conducted using FLAME in FSL with a significance threshold at a cluster corrected (*p* < 0.05, *Z* > 2.8) level. FreeSurfer was used to project the results onto a surface-rendered macaque template brain^62^. Table 1 shows the anatomical regions with significant electrically-induced esfMRI activity (x, y, z in the macaque atlas brain space) resulting from Site 1 or 2 stimulation. The contrast between Site 1 and Site 2 did not result in any cluster corrected (*p* < 0.05) voxels.

*Region of Interest analyses:* ROI analyses used anatomically defined regions from the macaque atlas^61^ registered to the macaque template brain^60^ and to each animal’s dataset. For vlPFC subregion analyses, we used anatomically delineated areas 44 and 45 from the atlas and FOP from prior work^36^. The MTL subregion analyses used anatomical regions corresponding to the PHG, entorhinal cortex (EC), subiculum (Sub), dentate gyrus (DG) and the CA1, 3 and 4 subregions (CA2 was not available in the human brain atlas). No voxels overlapped between ROIs. Polar plots using the ROIs (Fig. 4) show the average positive BOLD peak *Z*-value across the scanning runs.

*Statistical tests:* Mixed-design ANOVA models were used to examine ROI effects. The statistical test was implemented in SPSS 24 (IBM Corp, USA) and used scanning run ROI peak *Z*-values as the dependent variable, with between-subject factors of Monkey (M1, M2) and Species (in the cross-species comparison: Macaque), within-subjects factors of ROI (vlPFC or MTL ROIs), Hemisphere (left or right) and Stimulation site (Site 1 or 2) as covariate. We ensured that the data fit normality and equality of variance assumptions by using rank-based normalization and reporting Greenhouse-Geisser corrected results as required.

### Human subjects

Patients with intractable epilepsy requiring chronic invasive monitoring as part of their clinical treatment participated: *N* = 29; male = 19, female = 10; age range: 13 - 59 years old; median age = 34 years; handedness: right dominant = 21, left = 6, mixed = 2. All experimental procedures were approved by university IRB and written informed consent was obtained from all subjects. Depth and subdural surface platinum electrodes (Ad-Tech Medical Instruments) had been implanted for clinical monitoring. The patients’ demographic information and treatment outcomes are shown in Tables 3-4.

**Table 4.** Human patient clinical observations and surgical resection sites

| Patient ID | Clinical MRI finding | Surgery | Seizure onset zone |
| --- | --- | --- | --- |
| 292 | Left frontal focal encephalomalacia | left ATL+ left frontal seizure focus resection | Left mesial temporal lobe + left frontal lobe |
| 302 | Left MTS | Left ATL | Left mesial temporal lobe |
| 307 | Left insular cavernoma | Cavernoma resection | Left insula |
| 314 | Left occipital cortical dysplasia | No resection | Bilateral mesial temporal lobe |
| 316 | Right MTS | Right ATL | Right mesial temporal lobe |
| 320 | Right MTS | Right ATL | Right hippocampus |
| 330 | Right occipital cortical dysplasia | Right occipital lesionectomy | Left occipital lobe |
| 331 | Right MTS | No resection | Left mesial temporal lobe |
| 334 | Right temporal ganglioglioma | Right ATL | Right temporal pole, ventral surface of temporal lobe |
| 335 | Left MTS | No resection | Bilateral medial temporal lobe |
| 339 | Normal | No resection | Not determined |
| 352 | Left frontal cystic lesion | Left frontal regionectomy | Left frontal cystic mass |
| 357 | Normal | Left ATL | Left mesial temporal lobe |
| 369 | Right basal ganglia venous anomaly | Right ATL | Right mesial temporal lobe |
| 372 | Normal | Left ATL | Left temporal pole |
| 376 | Normal | Right ATL | Right mesial temporal lobe |
| 384 | Normal | Right ATL, Right frontal pole resection | Right mesial temporal lobe, Right frontal pole |
| 394 | Right temporal lobe cavernoma | Right ATL | Right amygdala |
| 395 | Left frontal lobe cavernoma | Left mesial frontal lobe resection including left frontal cavernoma | Left frontal lobe |
| 399 | Normal | Right ATL+ right ventral frontal seizure focus resection | Right mesial temporal lobe + possible right basal frontal lobe |
| 400 | Left MTS | Left ATL | Left mesial temporal lobe |
| 403 | Normal | Left ATL | Left mesial temporal lobe |
| 405 | Left frontal encephalomalacia | Left frontal regionectomy | Left frontal encephalomalacia |
| 413 | Normal | Right ATL | Right mesial temporal lobe |
| 418 | Right inferior temporal dysplasia | Right ATL | Right mesial temporal lobe |
| 423 | Normal | Left ATL | Left mesial temporal lobe |
| 427 | Right frontal encephalomalacia | Right frontal lobectomy | Right frontal lobe |
| 429 | Normal | Left ATL | Left mesial temporal lobe |
| 457 | Normal | Left ATL | Left mesial temporal lobe |
| 460 | Normal | Left ATL | Left mesial temporal lobe |

MTS: Mesial temporal sclerosis
ATL: Anterior temporal lobectomy

### Human esfMRI procedure

Details of the human esfMRI procedure and safety testing are available elsewhere^26^. In this study, we focused on data from auditory cortex stimulation with combined fMRI. Pre-electrode clinical implantation T1-weighted and T2-weighted structural MRI images were obtained at 3 Tesla (GE Discovery 750W scanner, 32 channel head coil): T1w inversion recovery fast spoiled gradient recalled (BRAVO) sequence, 1.0 × 1.0 × 0.8 mm^3^ voxel size, TE = 3.28 ms, TR = 8.49 ms, TI = 450 ms, FOV = 240 mm^3^, flip angle = 12 degrees; T2w: 3-D fast spin-echo (CUBE) sequence, 1.0 × 1.0 × 1.0 mm^3^ voxel size, TE = 77.21 ms, TR = 3200 ms, TI = 450 ms, FOV = 256 mm^3^, flip angle = 90 degrees.

During esfMRI scanning sessions, structural T1-weighted images were obtained using a Siemens Skyra 3 Tesla scanner (MPRAGE sequence with 1.0 × 1.0 × 1.0 mm^3^ resolution, TE = 3.34 ms, TR = 2530 ms, TI = 1100 ms, FOV = 256 mm^3^). A transmit and receive head coil was used for esfMRI sessions and for the session structural and functional scans. Gradient-echo, echo-planar imaging (GE-EPI) was used to obtain the T2* weighted BOLD scans (TR = 3.0 s, TE = 30 ms, slice thickness = 3.0 mm, FOV = 220 mm^3^, flip angle = 90 degrees, matrix size 68 mm^3^).

Stimulus isolators were used for electrical stimulation, connected to two of the available intracranial electrode contacts. Stimulus waveforms were computer controlled and electrical stimulation was induced via an optically isolated stimulation unit (AM Systems, Model 2200). The control computer received and timed the electrical stimulation via a trigger from the scanner indicating the start of each EPI volume acquisition. Stimulation was bipolar using adjacent contacts (inter-contact distance was 5 or 10 mm) with stimulus intensity between 9-12 mA using constant-current electrical stimulation. In-vivo impedance of the electrode contacts ranged from 1.5 kΩ to 5.5 kΩ at 100 Hz for both depth (cylindrical) and surface (disk) electrode contacts. Electrical stimulus waveforms were charge-balanced biphasic square waves (0.2 and 0.6 ms duration) with a 0.2 ms inter-stimulation pulse period at 100 Hz repetition as illustrated in Fig. 2B. Stimulation was delivered in blocks of 7 or 9 pulses repeated for 10 consecutive TRs followed by a 30 s no-stimulation baseline period. Each scanning run contained 10 stimulation and 11 no-stimulation blocks. Overall, 42 esfMRI runs in 19 patients were available for this study (Table 3). Stimulated sites in all of the individual brain are shown in Suppl. Fig. S2.

*Stimulation site categorization procedure:* The stimulation sites in the human esfMRI (and the electrical tractography, see below) were divided into two categories: 1) postero-medial HG sites (medHG); and 2) antero-lateral HG sites which included some sites on the planum temporale (latHG+PT). The number of esfMRI runs with Site 1 stimulation and Site 2 stimulation were 23 and 19, respectively.

The two sites were categorized according to the electrophysiological responses to click sounds presented at different rates^42^. Click trains of various repetition rates (0.2 ms square pulse, 25, 50, 100, 150 and 200 Hz, 50 presentations for each condition) were presented to the subject through earphones (ER4B Etymotic Research) binaurally fitted in custom-made ear-molds. The intracranial neurophysiological signal was recorded with an ATLAS system (NeuraLynx) at a sampling rate of 2 kHz (0.1 - 500 Hz acquisition filter). Raw (wideband) averaged potentials were calculated. The wideband averaged potentials were subsequently bandpass filtered centered at the click repetition rate (Windowed FIR filter with tap length of 250 sampling points, passband width 8 Hz). If averaged evoked potentials in response to click trains showed short-latency (<20 ms) waveform components and a frequency-following response to the 50 Hz or higher click rate, that contact was categorized as Site 1 (medHG) since the typical distribution of these sites are typically in the posterior to medial part of HG^42^. If the click-train induced averaged potentials showed clear wideband auditory evoked potentials but failed to show a strong frequency-following response, that site was categorized as Site 2 (latHG+PT); these types of responses are more typical in higher-order auditory regions including the antero-lateral parts of HG and planum temporale, see Suppl. Fig. S3.

*Electrode localization procedure:* The location of the implanted electrodes was determined by comparing pre- and post-electrode implantation structural T1w MRI scans. To compensate for potentially significant displacement of the electrode due to postoperative brain shift, post-implantation volumes were non-linearly warped into pre-implantation MRI volume space using a thin-plate spline (TPS) procedure with manually selected control points for the electrodes in three-dimensional space^26,63^. Between 50 and 100 control points throughout the brain are typically selected in this step.

Contact coordinates in the subject’s original space were transformed to standard MNI space using affine transformation and surface-based non-linear transformation implemented in FreeSurfer^62^.

For surface grid electrodes, the locations of all grid contacts were identified in a postoperative CT scan. This was accomplished by manually identifying the location of a subset of contacts in the grid on the basis of the characteristic hyper-intense radiological artifacts. Identified contacts included the depth electrode contacts or the full 64- or 96-grid fitted to these locations by TPS warping, using a negligibly small regularization parameter. Applying TPS allowed the non-linear deformation of the grid to be approximated. Accuracy of fitting was evaluated by visually comparing fitted contact locations with the contact artifacts in the CT and by verifying that inter-contact spacing fell within 0.2 mm of the expected 5-10 mm contact spacing.

After the initial grid locations were determined using CT, these were further corrected using a pre-explantation MR scan. Because displacement of brain parenchyma related to electrode mass-effect and post-operative swelling is often difficult to evaluate accurately on the CT scan, the results of CT-based localization were compared against a T1w MR scan obtained shortly before explantation. When significant discrepancy (greater than approximately 2 mm) was observed between CT-derived contact locations and corresponding magnetic susceptibility artifacts in the MR scan, a rigid linear transform was used to adjust grid positioning on the basis of clearly identifiable electrode-related artifacts in the MRI scan. The corner contacts were used as control points in this transformation. Individual T1w structural volumes were warped onto the CIT168 template brain (registered in MNI space, https://osf.io/hksa6/) with ANTs symmetric normalization algorithm^64,65^ and the contact coordinates in the original space warped onto the CIT168 template yielded the MNI coordinates used in this study.

### Human esfMRI data processing

The anatomical and functional imaging data were pre-processed using the fMRIPrep pipeline^66^.

### Anatomical image preprocessing

The high resolution T1w image was corrected for intensity non-uniformity using ‘N4BiasFieldCorrection’ and was used as a reference image throughout the workflow. The T1w-reference was then skull-stripped using ‘antsBrainExtraction.sh’ (ANTs 2.2.0) using the OASIS template as a target^64^. Spatial normalization to the ICBM 152 Nonlinear Asymmetrical template version (2009c) was performed through nonlinear registration with ‘antsRegistration’, using brain-extracted versions of both the T1w and template brains. Brain tissue segmentation of cerebrospinal fluid (CSF), white-matter (WM) and gray-matter (GM) was performed on the brain-extracted T1w using FAST in FSL.

The subject’s pre-electrode implantation structural MRI and the template brain (MNI-152-NonLinear-2009c Asymmetrical brain) were processed with FreeSurfer ‘recon-all’ procedure to create the surface mesh. For improved pial surface reconstruction, cortical parcellation was facilitated via the T2w structural scans (1.0 × 1.0 × 1.0 mm^3^) obtained during the same imaging session whenever possible. For mapping the BOLD and electrophysiological response onto the brain surface, the FreeSurfer surface meshes were further processed with AFNI’s @SUMA_Make_Spec_FS to create standard icosahedron surfaces with various mesh densities.

### Functional data preprocessing

For each of the esfMRI runs per subject, first a reference volume and its skull-stripped version were generated using fMRIPrep. The T2*-weighted reference was then co-registered to the T1w reference using FLIRT in FSL with the boundary-based registration cost-function. Co-registration was configured with nine degrees of freedom to account for T2*w distortions. Head-motion parameters with respect to the BOLD reference (transformation matrices, and the six rotation and translation parameters) were estimated before any spatiotemporal filtering using FLIRT in FSL. EPI scans were slice-time corrected using ‘3dTshift’ from AFNI^64^. The BOLD time-series (including slice-timing correction when applied) were resampled onto their original, native space by applying a single, composite transform to correct for head-motion and susceptibility distortions. The BOLD time-series were then resampled to the MNI-152-NonLinear-2009cAsymmetrical standard space. Several confounding time-series were calculated based on the preprocessed BOLD: framewise displacement (FD), DVARS and three region-wise global signals. FD and DVARS are calculated for each functional run, both using the implementation in Nipype @power_fd_dvars.

The three global signals, CSF, WM and whole-brain masks were extracted, though not used as nuisance regressors. Additionally, a set of physiological regressors were extracted to allow for component-based noise correction (CompCor). Principal components were estimated after high-pass filtering the preprocessed BOLD time-series (using a discrete cosine filter with 128 s cut-off). We also defined two CompCor variants: temporal (tCompCor) and anatomical (aCompCor). Six tCompCor components were also calculated from the top 5% voxel variability within a mask covering subcortical regions. This subcortical mask is obtained by heavily eroding the brain mask, which ensures it does not include cortical gray matter. The head-motion estimates calculated in the motion correction step were also added to the confounding variables file. All resampling was then performed using a single interpolation step by composing all the pertinent transformations (i.e., head-motion transform matrices, susceptibility distortion correction, when available, and co-registration information to anatomical and template spaces). Gridded volumetric resampling was performed using ‘antsApplyTransforms’ (ANTs), configured with Lanczos interpolation to minimize smoothing effects (@lanczos).

The above fMRIPrep processing pipeline generally yielded good registration between anatomical and functional imaging data, even with signal dropout due to the intracranial electrodes. If misalignment was obvious in the visual inspection of EPI to T1w registration or as reported in fMRIPrep, AFNI’s coregistration program discarded parts of the functional data that had significant signal dropout. Here, the functional data was clipped in the sagittal plane and this volume was used for finding the coregistration parameters using AFNI’s ‘align_epi_anat.py’ program with normalized mutual information as a cost function. During this step, the part of the brain contaminated with the intracranial electrodes was not used for coregistration.

### General Linear Modelling

The preprocessed functional datasets were subjected to univariate general linear model (GLM) analysis using AFNI’s 3dDeconvolve routine. For the GLM analysis, functional data is spatially smoothed with a Gaussian kernel (FWHM = 6.0 mm). Stimulus times were convolved with a 1-parameter gamma function. Baseline detrending was applied with a Legendre polynomial (5 degrees). Volumes (TRs) that showed large levels of motion (FD > 1.0 mm) and adjacent TRs were discarded. The six tCompCor components extracted above and the FD time-series were added to the baseline model as nuisance regressors.

A brain mask was created before spatial smoothing using intensity thresholded EPIs excluding areas of signal dropout from the electrodes contributing to the fMRI analyses. The clinically determined seizure onset zone (SOZ) was also excluded from the brain mask. If the SOZ affected any part of an ROI, the result for that run and ROI was excluded from further analysis. We show an incidence map in Suppl. Fig. S3 showing the data across the brain that were included.

For the higher-level group analysis we used AFNI’s ‘3dREMLfit’^64^. Datasets showing evidence of a response anywhere within the brain mask (false-discovery rate corrected, Z >2) were submitted to higher-level analysis. Resulting statistical maps were subjected to multi-level mixed-effects analysis using ‘3dMEMA’. The first- and higher-level GLMs were conducted in standard space (MNI-152-NonLinear-2009c Asymmetrical).

### ROI analyses

Regions of interest analyses of the vlPFC and MTL used anatomically defined ROIs from standard anatomical atlases of the human brain. The vlPFC subregion analyses included as ROIs areas 44 and 45, and the frontal operculum (FOP). For area 44 and 45, the parcellation is based on the Jülich histological (cyto- and myelo-architectonic) atlas using a 25% probability threshold^67^. The human FOP ROI is based on prior work^36^. The MTL subregion analyses used anatomical regions corresponding to the following subregions: subiculum (Sub), dentate gyrus (DG) and the CA1, 3 and 4 subregions, using FreeSurfer’s hippocampal subfield segmentation (v6.0)^68^. For the entorhinal cortex (EC) and parahippocampal gyrus (PHG), we used FreeSurfer’s cortical segmentation (aparc+aseg files) from the Desikan-Killiany atlas. No ROIs had overlapping voxels. Polar plots (Fig. 4) show the average positive BOLD peak *Z*-value, with variability across the scanning runs in the humans.

### Statistical tests

Mixed-design ANOVA models were used to examine ROI effects. The statistical test used scanning run ROI peak *Z*-values as the dependent variable, with between-subject factors of Human and Species (in the cross-species comparison: Human), within-subjects factors of ROI (vlPFC or MTL ROIs), Hemisphere of activation (left or right) and Stimulation site (Site 1 or 2) and Stimulated hemisphere (left or right) as covariates. We ensured that the data fit normality and equality of variance assumptions of the models or transformed the data to achieve normal distribution and report Greenhouse-Geisser corrected results, as required.

### Electrical stimulation tract-tracing (esT)

Electrical stimulation neurophysiological tractography (esT) was conducted in human patients (*N* = 13, Table 3) according to general methods described previously^16,28^. Here, we used a single constant current electrical stimulation pulse (biphasic charge-balanced square wave, duration = 0.2 ms/phase, 9 or 12 mA). The electrical pulses were delivered through stimulus isolators connected to the intracranial electrodes in a bipolar configuration (always connected to adjacent contacts). The inter-stimulus interval was set to 2 s and repeated 60 times. The intracranial EEG signal was recorded using the ATLAS system (NeuraLynx) with a sampling frequency of 8 kHz. A disk contact in the subgaleal space was used as a reference electrode for the recordings. Stimulated sites included HG, STG, vlPFC and hippocampal contacts. Average potentials were calculated for each contact after rejecting trials that contained large amplitude non-physiological signals after applying a high-pass filter (4th order Chebyshev type 2, −6 dB roll-off at 3 Hz) and de-meaning the potentials. The trial exclusion criterion for rejection was a signal greater than 3 times the interquartile range above the 75th percentile of the amplitude distribution.

### Spline-Laplacian correction

A Laplacian procedure was applied to reduce non-specific neurophysiological or electrical stimulation effects evident as cross-correlated potentials common to many electrodes from a common source or far-field potentials^69,70^. This procedure spatially corrects the average potentials after spline interpolation. The Laplacian operation is a spatial high-pass filter (second derivative) and the resulting potentials are reference independent and de-emphasize far-field effects or those from volume conduction. For depth electrode potentials, a 1-dimensional spline-Laplacian was calculated using the inter-contact distance information to calculate a spherical spline-laplacian^71-73^. For potentials from surface grids and strip electrodes, we used the spherical coordinates corresponding to each contact’s MNI coordinates based on FreeSurfer’s spherical surface mapping^62^. The regularization parameter for spherical surface spline-Laplacian computation was determined by generalized cross-validation^72^ and the spline-flexibility parameter was set to 3. The magnitude of esT responses was quantified by comparing the root mean square (RMS) values of the Laplacian transformed waveforms between the post-stimulation period (10 - 200 ms after stimulation onset) and pre-stimulation period (−500 to −10 ms) prior to the electrical stimulation pulse.

### esT movie creation method

To make the movies, continuous intracranial recordings were cut into trials (−1 to 1 s around electrical stimulation). The following procedure that we used yields similar results to one extracting high-frequency power (gamma and high-gamma band), but it avoids the need for temporal band-pass filtering and frequency decomposition. Stimulation artifacts (−5 to 5 ms from stimulation) were first replaced with the time-reversed waveform of the same length (10 ms) during an artifact free pre-stimulus period (−10 to −5 ms). Then a peri-stimulus waveform of 12 ms duration (−6 to 6 ms) was smoothed twice with median filters (length 3 ms, followed by 5 ms). This process was done for all single-trial waveforms. The trials were then re-referenced to the average of the surface grid contacts, to reduce the volume conducted artifact waveforms common across the recording surface grid. To extract induced responses (not phase-locked components), each channel’s averaged potential in the re-referenced signal was subtracted from the single trials in that channel^74^. Induced response magnitude was calculated by taking the trial average of the full-wave rectified signal.

The magnitude of stimulation-induced responses was calculated by taking the logarithm of the ratio of the magnitude in the pre-stimulus period (0.5 to 0.3 s before stimulus onset) and post-stimulus period. This yields the relative magnitude change with respect to the pre-stimulus period in dB. To make the movies, averaged induced responses of the surface contacts were calculated for non-overlapping temporal windows (length 5 ms), then color-coded and plotted onto the MNI template brain for each experimental run and temporal window. We employed bootstrapping (1000 iterations) of the mean activity within 5 ms temporal windows from the pre-stimulus period (55 to 7.5 ms before stimulation onset) and thresholded the response at the lower and upper 2.5 % points. Responses not exceeding this threshold were set to 0 and thus were not mapped to the brain surface. For patients who had multiple esT sessions for either medHG or latHG sites, we averaged responses for each site separately to create the movies (Suppl. Movies 1-6).

### Speech sound recording experiment and Granger Causality analyses.

To examine the brain’s effective connectivity under a natural sensory stimulation setting, we examined neurophysiological responses to speech sounds. Many of the subjects were the same neurosurgical patients (*N* = 8) who took part in the electrical tractography study (Table 3-4).

### Speech stimuli

The experiment used a speech presentation paradigm previously described^75,76^. The speech sounds were common monosyllabic consonant-vowel-consonant English words, e.g., “cat”, “dog”. All sounds were normalized to the same root mean square amplitude and edited to be 300 ms in duration with 5 ms amplitude rise and fall times. Sounds were delivered binaurally via insert earphones (ER4B, Etymotic Research, Elk Grove Village, IL, USA) integrated into custom-fit ear molds. Sound delivery was controlled using Presentation software (Version 16.5 Neurobehavioral Systems). Altogether 80 presentation trials were presented during two experimental blocks. The subjects were asked to press a button when they heard a target word (only non-target word responses analyzed here) using their index fingers ipsilateral to the hemisphere from which the recordings were made.

### Event-related spectral analysis of LFP

Intracranial recording data were downsampled to 1000 Hz. The analysis of the neurophysiological responses focused on five frequency bands: theta (4-8 Hz), alpha (8-14 Hz), beta (14-30 Hz), gamma (30-70 Hz) and high gamma (70-150 Hz) denoised using a demodulated band transform-based algorithm^77^. Event-related spectral perturbations were calculated by log-transforming the power for each center frequency and normalizing it to the baseline (mean power in the pre-stimulus reference interval of −200 ms to −100 ms before stimulus onset). The waveforms were then averaged across trials.

### State-space Conditional Granger Causality (CGC) analysis

CGC was used to investigate the directional influence between brain regions during speech processing. The method is multivariate and conditional, in the sense that simultaneous time series from a collection of electrodes are included in order to account for both direct and indirect influences between contacts. Intracranial recordings were downsampled to 100 Hz and sectioned into trials from −200 to 1000 ms relative to sound onset. Intuitively, CGC tests if activity in a source area can be used to predict subsequent activity on a target area. We estimated spectral CGC in 500 ms sliding windows in steps of 50ms to construct trial-averaged time-frequency CGC representations between selected pairs of electrodes^78^. Prior to time-frequency CGC analysis, the mean at individual time points across trials was subtracted from the single trial responses and then scaled by the standard deviation.

Several significant problems arise in applying standard Vector Auto-Regressive (VAR) based CGC models to intracranial recordings, which are related to downsampling and nonstationarities^79^. Most of these recorded time-series contain a moving-average (MA) component that may not be adequately modeled using VAR due to the intractably large model order necessary to handle the MA component. It has been shown that spectral CGC estimates can be obtained with high computational reliability using estimation approaches based on single model-fits. The state-space model addresses a number of theoretical and practical problems related to spectral CGC estimation, e.g., see^80^. Spectral CGC was directly computed using Geweke’s formulations based on the estimated state-space innovations covariance matrix^81,82^, cross-spectral densities and transfer functions^83,84^.

State-space models use state variables to describe a system by a set of first-order differential or difference equations, rather than by one or more VAR *n*th-order differential or difference equations. State variables can be reconstructed from the measured ecordings but are not themselves measured during an experiment. For modeling directional influence in the brain, it is possible to directly express the interactions between different regional signal time series as a state-space model defined by:

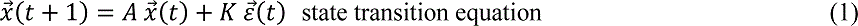

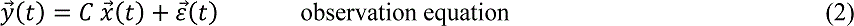

where x^⃗^(t) is an unobserved (latent) *m*-dimensional state vector, and ε^⃗^(t) is the vector of the innovations or prediction errors. The observed vector of time-series y^⃗^(t) corresponds to the recordings from regions in the targeted network. The state transition matrix *A*, observation matrix *C* and the steady-state Kalman gain matrix *K* are estimated using a subspace method^85^. Subspace methods are optimal for state-space model parameter estimation, especially for high-order multivariable systems^86^. The order of the state-space model was 25, which corresponds to the vector size of x⃗(t).

To statistically evaluate the reliability of the connectivity results we used a phase-randomization surrogate data technique to construct a null distribution^87,88^. This method consists of randomly shuffling the Fourier phases of each of the intracranial recordings, which generates uncorrelated data with preserved autocorrelation properties. For each original data set, 500 surrogates were generated, and CGC values that exceeded the 95% threshold were unmasked in the time-frequency representations.

## EXTENDED DATA

Supplementary Movies M1-M6. Electrical tractography (esT) results: impact of Heschl’s Gyrus stimulation on high gamma responses in the brain, as a function of time. Induced high gamma power results from electrically stimulating contacts in medial or lateral Heschl’s gyrus are shown in three subjects.

- M1: Movie_1_AEP-Ave-medHG_movie_estt_423: Subject 423 stimulation of medHG
- M2: Movie_2_AEP-Ave-latHG_movie_estt_423: Subject 423 stimulation of latHG+PT
- M3: Movie_3_AEP-Ave-medHG_movie_estt_384: Subject 384 stimulation of medHG
- M4: Movie_4_AEP-Ave-latHG_movie_estt_384: Subject 384 stimulation of latHG+PT
- M5: Movie_5_AEP-Ave-medHG_movie_estt_372: Subject 372 stimulation of medHG
- M6: Movie_6_AEP-Ave-latHG_movie_estt_372: Subject 372 stimulation of latHG+PT

## SUPPLEMENTARY MATERIALS

**Supplementary Figure S1.**
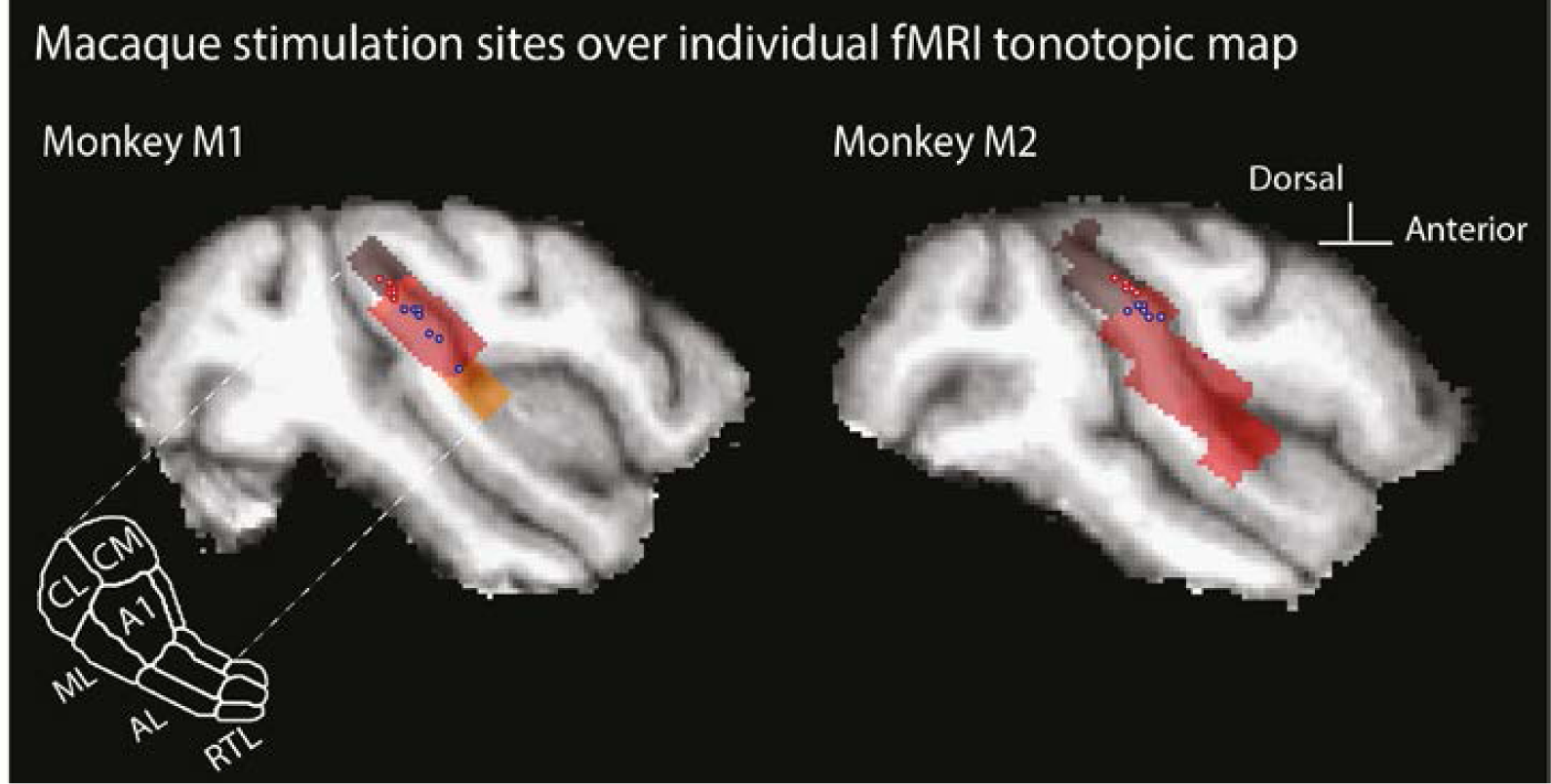
Macaque stimulation sites overlaid on individual animal fMRI tonotopic maps. Shows the stimulation sites (Site 1 blue circles; Site 2 red circles) in the two macaques with an overlaid individual animal fMRI tonotopical map (gray to orange shading).

**Supplementary Figure S2.**
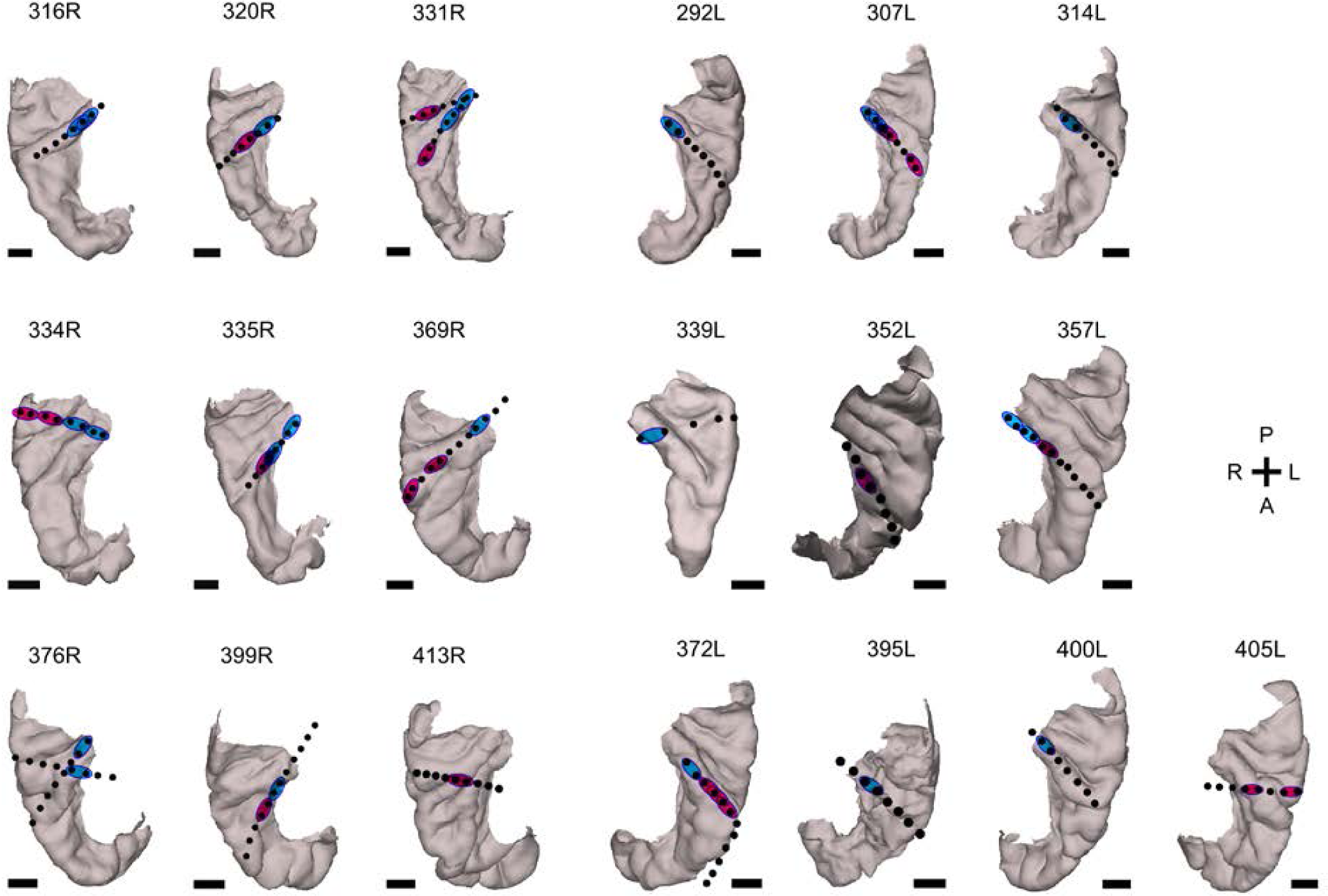
Individual human depth electrode contact locations for esfMRI. Shown are views looking down on the superior temporal plane. Black circles identify the clinically placed contacts and the blue/red regions the two contacts used for bipolar stimulation, respectively in the medHG or latHG+PT sites, shown on each individual’s anatomy. Whenever more than two contacts are shown in red is an indication of other pairs of adjacent contacts that were stimulated in a separate testing run. Scale bars 10 mm.

**Supplementary Figure S3.**
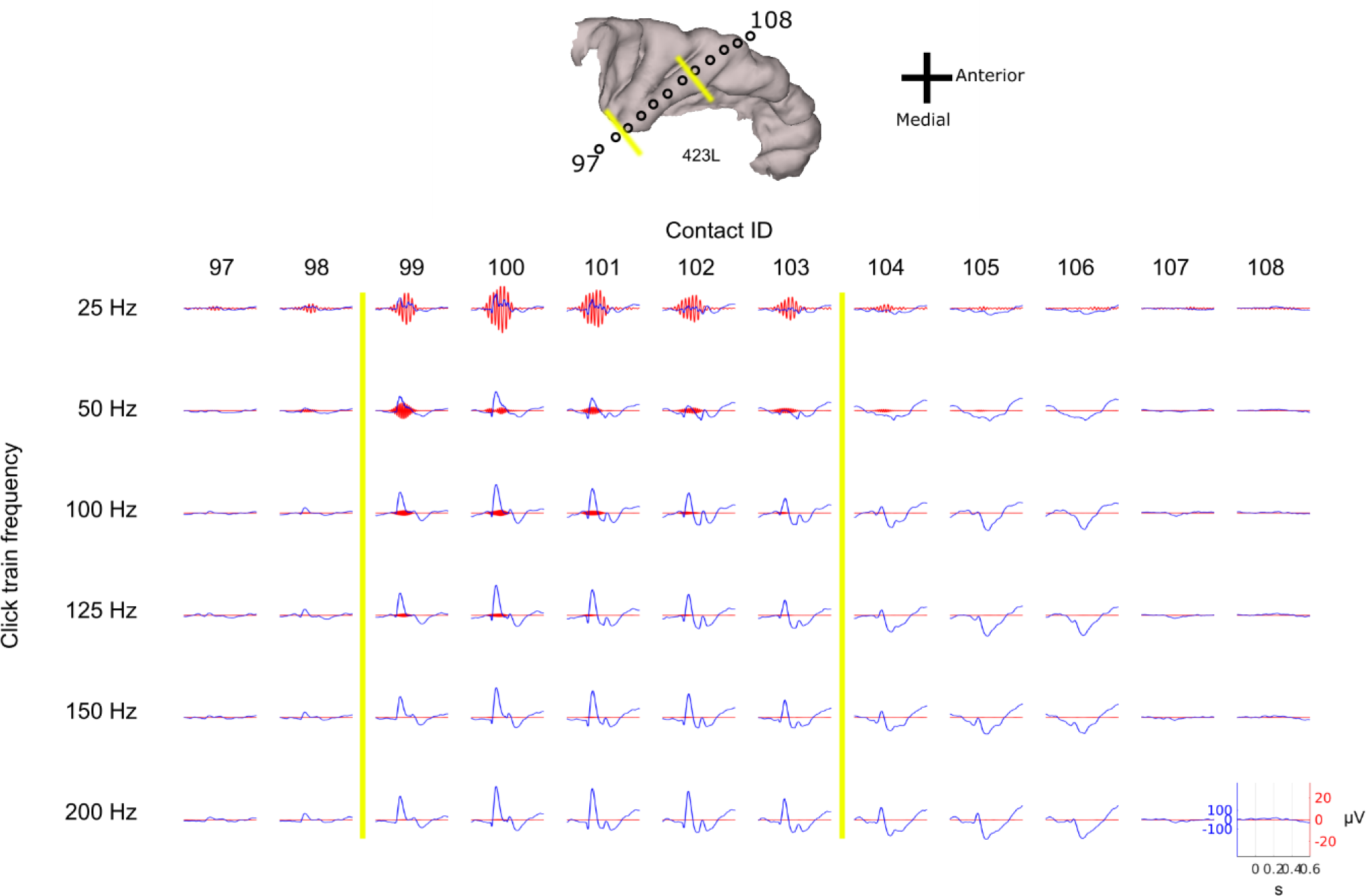
Click frequency following response used to identify Site 1 (medHG) sites in humans. Shown is how the click evoked neurophysiological response was used to identify medial HG contacts by their click-frequency following response. In this example, traces in blue show the auditory evoked potentials. Traces in red are the high-pass filtered frequency following response. Contacts between the two yellow lines show a strong frequency following response to 25-100Hz and were thus assigned to medHG. Contacts on the right are assigned to Site 2 (latHG+PT) because they respond to the sounds but do not show a clear high frequency following response. The non-responsive contacts at the flanks of these were not used.

**Supplementary Figure S4.**
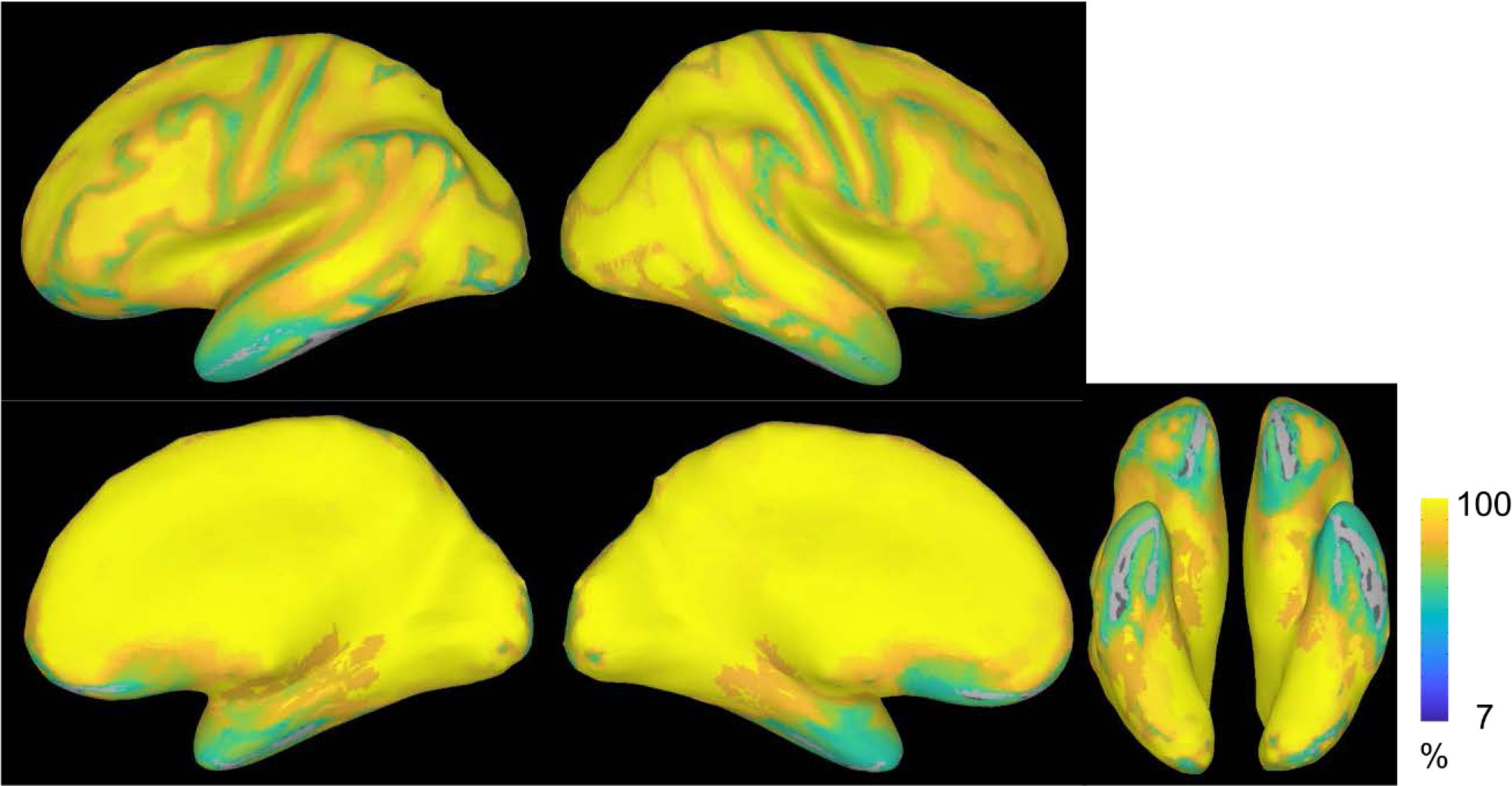
Human electrode contact locations affecting fMRI signal: preserved fMRI signal map. Regions where the fMRI signal was contaminated by the electrodes were masked and excluded from analysis. We also excluded regions that were resected, typically in the anterior medial temporal lobe (Table 4). Here is show an incidence map identifying the regions most affected by signal drop out (blue and green colored regions), which includes areas where the fMRI signal is also affected by sinuses (orbitofrontal cortex). Orange to yellow color shows the percentage of human esfMRI runs that were available for analysis (not masked by intracranial electrodes, weak signal or outside of the epileptic foci regions).

**Supplementary Figure S5.**
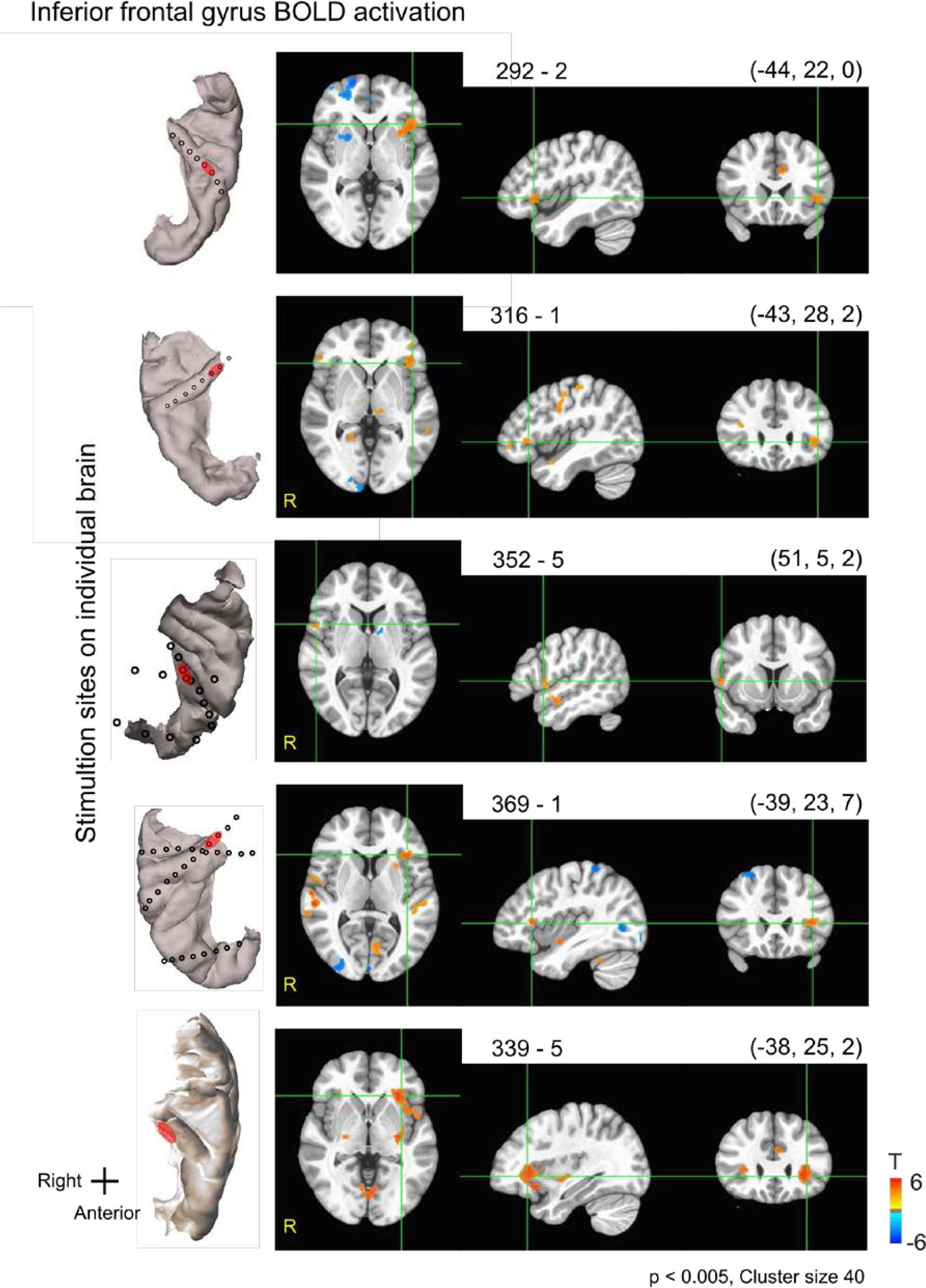
Individual human es-fMRI activation results involving vlPFC. Shown are several individual human results showing substantial vlPFC (inferior frontal cortex/gyrus) activation p < 0.005, cluster size = 40. Numbers above the panels indicate the subject number and testing run, followed by MNI x, y, z coordinates at the cross-hairs.

**Supplementary Figure S6.**
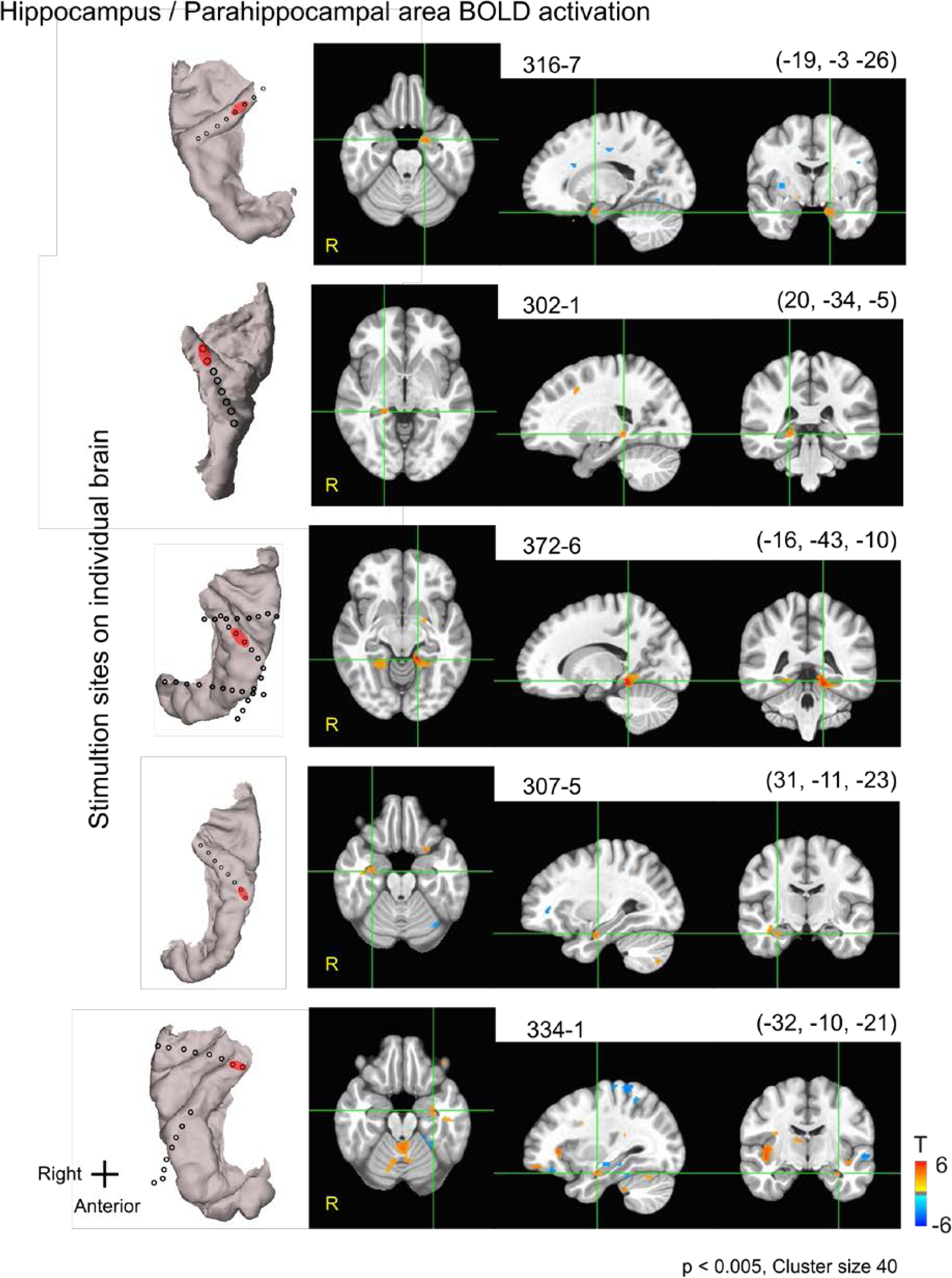
Individual human es-fMRI results involving medial temporal lobe (MTL). Shown are several individual human results showing substantial MTL activation, p < 0.005, cluster size = 40. Format as in Suppl. Fig. S5.

**Supplementary Figure S7.**
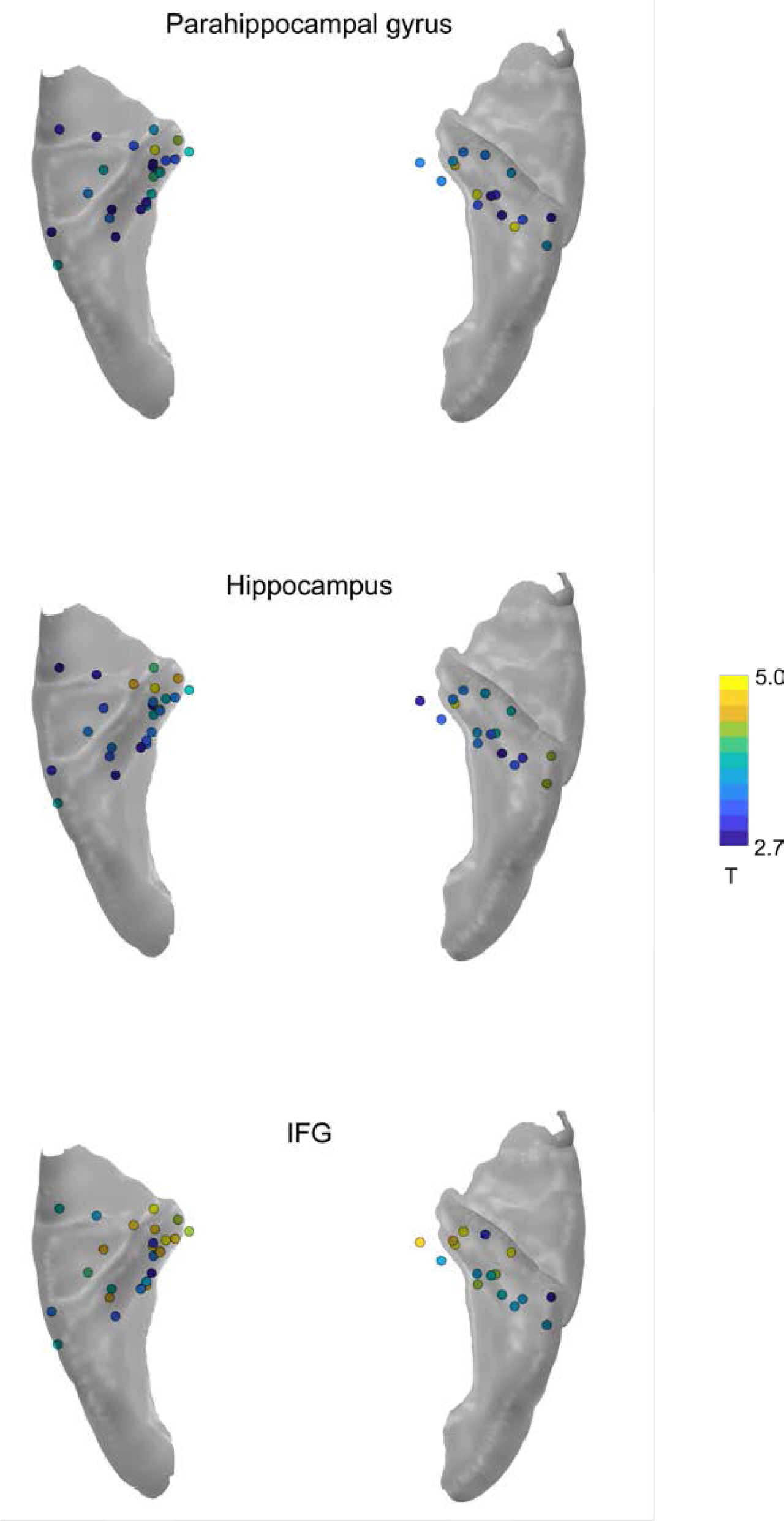
Impact of stimulation sites on the MTL and vlPFC contact sites. Shown are the auditory cortex regions that when stimulated resulted in the displayed strength of fMRI responses on vlPFC (inferior frontal gyrus: IFG) and MTL regions. Auditory cortex (Heschl’s gyrus) stimulation sites are color coded from blue to yellow color map according to the strength of the fMRI BOLD response elicited in the parahippocampal, hippocampal, or vlPFC (IFG) regions. Although stronger responses tend to be seen from the more medial HG contacts, there are also contacts bordering other regions that also elicit strong responses and passive current spread is a factor, see manuscript text discussion.

**Supplementary Figure S8.**
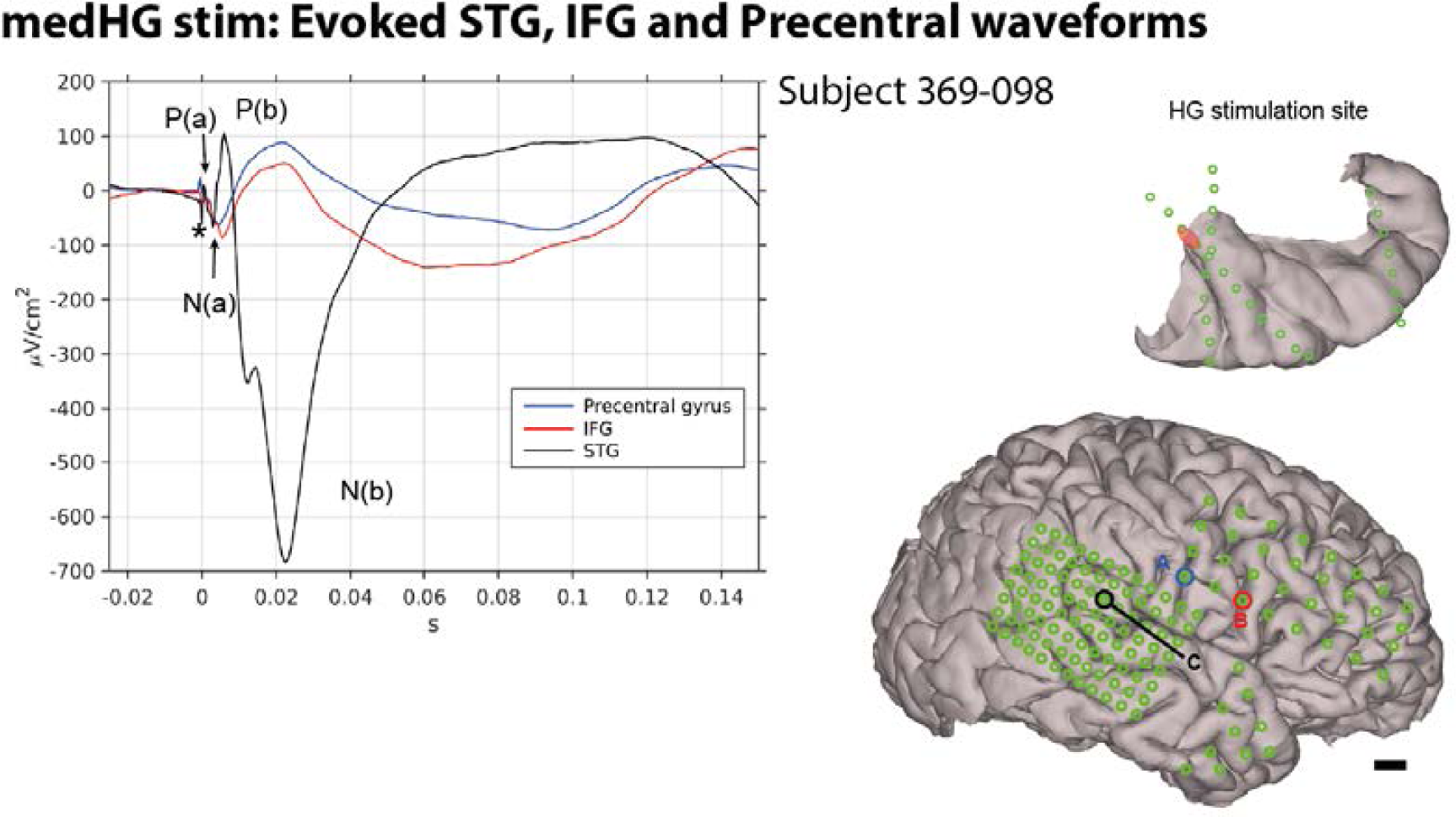
Impact of medHG stimulation on STG evoked responses in subject 369. The early positivity P(a) and negativity N(a) in this subject at least qualitatively replicate the effects reported previously of medHG stimulation impact on the latency of the recorded potentials in the STG (Brugge et al., 2003). Upper right image shows the adjacent stimulation contacts in HG (shaded red area). Lower right shows the recording contact electrode locations and potentials evoked in inferior frontal gyrus (IFG) within vlPFC and the precentral gyrus, see manuscript text. Scale bar in bottom right of figure is 8 mm for top image and 1 cm for whole brain.

